# Cell-type specific gating of gene regulatory modules as a hallmark of early immune responses in Arabidopsis leaves

**DOI:** 10.1101/2025.08.01.668105

**Authors:** Shanshan Wang, Ilja Bezrukov, Pin-Jou Wu, Hannah Gauß, Marja Timmermans, Detlef Weigel

## Abstract

In plants, many cell types contribute to immunity, but what division of labor exists among cell types when immunity is activated? We compared, at single-cell resolution, the response of *Arabidopsis thaliana* leaf cells during pattern-triggered and effector-triggered immunity (PTI/ETI), sampled at 3 and 5 hours after infection with *Pseudomonas syringae* DC3000. Core defense modules were broadly shared across cell clusters, but their activation varied in timing and intensity, with key immune receptors also showing cell type–specific expression dynamics. We identified distinct mesophyll cell populations based on their resilience patterns: after the initial response, some cells continue to express defense genes at high levels during both PTI and ETI, while others quickly reinitiate growth-related gene expression programs, but only during PTI. Gene regulatory network inference revealed WRKY-regulated modules enriched in cells sensing effectors, while SA biosynthesis regulators were activated in complementary clusters. Finally, we used the *cue1*-6 mutant to demonstrate that core immune responses are robust despite altered leaf architecture. In addition, we uncovered cryptic defense pathways, including sucrose-responsive modules, in this mutant. By capturing early immune responses at high resolution, our study reveals cell type–specific coordination of plant immunity and provides a framework for decoding immune signaling networks.

## Introduction

Plant immunity operates through two interconnected layers: pattern-triggered immunity (PTI), initiated by cell-surface pattern recognition receptors (PRRs) that sense conserved microbial patterns, and effector-triggered immunity (ETI), a more robust response mediated by intracellular nucleotide-binding domain and leucine-rich repeat (NLR) proteins recognizing pathogen effectors (Jones & Dangl, 2006). While these layers are known to act synergistically and potentiate each other (Ngou *et al*., 2021; Yuan *et al*., 2021), how they are coordinated within the heterogeneous landscape of leaf tissues, especially at the earliest infection stages, remains incompletely understood.

Central to PTI are Receptor-Like Kinases (RLKs) and their close homologs, Receptor-Like Cytoplasmic Kinases (RLCKs), which form a large and diverse family of membrane-localized receptors that perceive pathogen-associated molecular patterns (PAMPs) and initiate downstream signaling cascades (Shiu *et al*., 2004; Boutrot & Zipfel, 2017; Ngou *et al*., 2022). RLKs thus often represent the first line of defense by detecting extracellular cues and triggering immune signaling. In parallel, NLR proteins, can be grouped into CNL (coiled-coil-NLR), TNL (TIR-NLR), RNL (*RPW8-like* coiled-coil-NLR) based on the structure of N-terminal domains, act as intracellular immune sensors that can detect specific pathogen effectors, activating robust ETI responses often associated with localized cell death and systemic resistance (Jones *et al*., 2016; Duxbury *et al*., 2021). Given their fundamental roles in pathogen perception and immune activation, understanding the spatial and temporal expression patterns of RLKs and NLRs across diverse leaf cell types is critical to deciphering how plants coordinate effective immunity at the cellular level.

Bulk RNA sequencing studies have revealed distinct transcriptional dynamics in PTI and ETI: While PTI induces rapid, transient changes in gene expression, ETI elicits more robust and sustained responses (Tsuda & Katagiri, 2010; Mine *et al*., 2018; Bjornson *et al*., 2021). PTI and ETI share a large cohort of induced genes (Navarro *et al*., 2004), even though there are differences in downstream regulatory mechanisms. While salicylic acid (SA), jasmonic acid (JA) and ethylene act primarily synergistically during PTI, they do so in a more redundant manner during ETI (Hillmer *et al*., 2017; Mine *et al*., 2018). Recent studies have also demonstrated mutual potentiation between PTI and ETI, where each pathway amplifies the other’s transcriptional output, with the PTI/ETI crosstalk enhancing defense durability (Ngou *et al*., 2021; Yuan *et al*., 2021).

The *Arabidopsis thaliana* (Arabidopsis) leaf is a mosaic of specialized cell types, with the major types being mesophyll, epidermal, guard, and vascular cells (Liu *et al*., 2020; Lopez-Anido *et al*., 2021; Kim *et al*., 2021). In addition, there are many transient cell states shaped by developmental gradients, metabolic activity, and microenvironmental cues (Lopez-Anido *et al*., 2021; Tenorio Berrío *et al*., 2022; Procko *et al*., 2022; Guo *et al*., 2025). Recent spatial transcriptomic and single-cell analyses have revealed that even within a single tissue type, such as the mesophyll, cells exhibit region-specific functional specialization. While photosynthesis is the primary function of mesophyll cells, gene expression profiling demonstrates that photosynthetic efficiency and related processes vary along the medio-lateral axis of the leaf (Xia *et al*., 2022). A comprehensive single-nucleus atlas of Arabidopsis has demonstrated that leaf cells differ markedly in their developmental age and metabolic status, with senescence and nutrient allocation occurring in a highly coordinated yet cell-type–specific manner across the leaf (Guo *et al*., 2025). Because of this extensive cellular and functional heterogeneity, leaves are poised to respond to pathogen challenges not uniformly, but with finely tuned, cell-type and context-dependent immune programs.

Recent advances in single-cell transcriptomic and epigenomic technologies have revolutionized our understanding of plant immunity, revealing in considerable detail temporal and spatial heterogeneity in immune responses. For example, single-cell RNA-seq (scRNA-seq) analyses of Arabidopsis plants infected with *Pseudomonas* highlighted both cell specialization and coordination of immune responses between cell types (Delannoy *et al*., 2023), as well as immune active cell states in mesophyll cells with elevated expression of *FRK1*, *CBP60G* and *LipoP1*, and particularly susceptible cell states marked by *EXPA10* and aquaporin genes (Zhu *et al*., 2023). Tang and colleagues (Tang *et al*., 2023), working with Arabidopsis and the fungal pathogen *Colletottrichum higginsianum*, found vasculature-specific enrichment of NLR expression and guard cell-specific regulation of abscisic acid (ABA) signalling, which mediates stomatal closure upon direct fungal contact. Single-nuclei RNA-seq (snRNA-seq) combined with single-cell ATAC-seq revealed rare PRIMER cells as immune signaling hubs, with high expression of *BON3* and genes whose promoters are enriched for CAMTA and GT-3A motifs, while adjacent bystander cells upregulate systemic resistance genes such as *ALD1* and *FMO1* (Nobori *et al*., 2025).

Here, we present a single-cell transcriptomic atlas of Arabidopsis wild-type and mutant leaves challenged with either virulent (“empty vector”, EV) or avirulent (expressing AvRpt2) *Pseudomonas syringae* pv. *tomato* (*Pst*) DC3000. DC3000 suppresses plant immunity using a suite of effector proteins and toxins, but when it also expresses the AvrRpt2 effector, which is recognized by the NLR receptor RPS2, plants mount an ETI response (Whalen *et al*., 1991; Kunkel *et al*., 1993; Mackey *et al*., 2003). We show that early immune responses are driven by a set of shared co-expression modules whose activation is gated at the level of cell types. Key regulons, controlled by GT-3a, and SA responsive or biosynthesis related TFs coordinate these modules within distinct mesophyll clusters. We also identify two contrasting mesophyll populations: one that always prioritizes defense over growth and another that does the opposite – but only during PTI. Moreover, we demonstrate that receptor expression dynamics further sculpt a heterogeneous yet robust immune response landscape. Finally, by profiling *cue1*-6 mutants, which have highly altered leaf architecture and impaired shikimate biosynthesis (Li *et al*., 1995; Streatfield *et al*., 1999; Voll *et al*., 2003; Pérez-Pérez *et al*., 2013), we demonstrate the robustness of immune modules in the face of major shifts in cell type populations in leaves and reveal a role for sucrose-derived pathways in bolstering early defense.

## Results

### A Single-Cell RNA-seq Atlas of Early Responses to Virulent and Avirulent *Pseudomonas* Infection

We generated a time series of scRNA-seq data for leaves of the Arabidopsis reference accession Col-0 and the *cue1*-6 mutant infiltrated with either virulent *Pst* DC3000 (EV) or avirulent *Pst* DC3000 (AvrRpt2), and MgCl_₂_ for mock samples (Fig. 1A). Leaves were syringe-infiltrated at a low dose of OD_₆₀₀_ = 0.005. At this dose, only some cells are expected to have direct contact with the pathogen during early stages of infection (Zhu *et al*., 2023; Delannoy *et al*., 2023; Nobori *et al*., 2025). We harvested the samples at 3 hours post infection (hpi) and 5 hpi to capture the earliest transcriptional changes. To minimize circadian effects, isolation of protoplasts and subsequent 10x Genomics droplet scRNA-seq preparation were always performed at the same time of day; infiltrations therefore were done at -5 h, -3 h, or -0 h relative to harvest and sample processing.

**Fig. 1.**
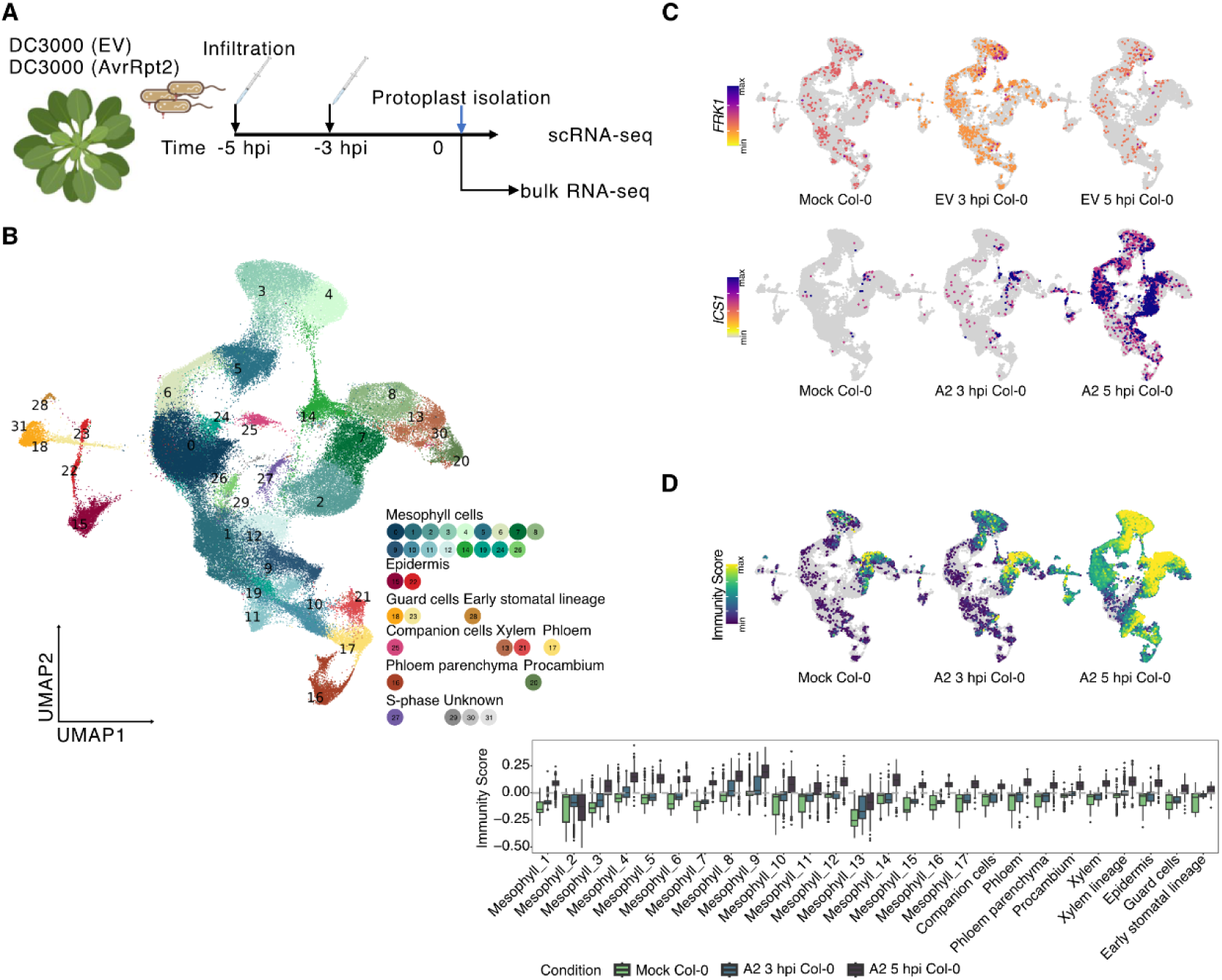
Single-cell RNA-seq atlas of Arabidopsis leaves during early *Pst* DC3000 infection. **A.** Experimental design. Fourteen-day-old rosettes were syringe-infiltrated with infiltration buffer only (mock), *Pst* DC3000 (empty vector, EV), or *Pst* DC3000 (AvrRpt2) at OD_₆₀₀_=0.005. Leaves were sampled 3 hours post-inoculation (hpi) and 5 hpi. Protoplasts were immediately isolated for single-cell RNA-seq (scRNA-seq) and bulk RNA-seq. **B.** Uniform manifold approximation and projection (UMAP) of transcriptomes from 109,192 cells, colored and numbered by Seurat cluster. Clusters were assigned to major cell types with known marker genes: mesophyll, epidermis (including guard-cell lineage), companion cells, phloem (parenchyma and procambium), xylem, and an S-phase cluster. **C.** Layered UMAP plots showing per-cell expression of the defense marker genes *FRK1* (top) and *ICS1* (bottom). Color scale denotes relative normalized transcript abundance. **D.** Top: UMAPs of the “Immunity Score”, computed with AddModule function in Seurat from bulk-RNA-derived DEGs in mock (green), A2 3 hpi (light teal), and A2 5 hpi samples (dark teal). Bottom: Box plot showing the distribution of Immunity Score in each cell cluster, comparing mock, A2 3 hpi and A2 5 hpi samples. EV and A2 are short for *Pst* DC3000 (EV) and *Pst* DC3000 (AvrRpt2), respectively.

After quality filtering, we retained data from 109,192 cells from 15 samples (mean ≈ 6,775 unique molecular identifiers (UMIs) and 2,150 detected genes per cell; see Methods, Fig. S1). Clustering and marker-based annotation with Seurat (Hao *et al*., 2021, 2024) identified 29 cell clusters that we could assign to the major histological cell types in the leaf, mesophyll (17 clusters), which constituted as expected the majority of cells; vascular tissue, including xylem (2 clusters), companion cells, phloem, phloem parenchyma, and procambium, epidermis (2 clusters), including non-guard cells (called “epidermis” from here on) along with guard cells (2 clusters) and cells belonging to the early stomatal lineage, as well as cells in S-phase (total 29 clusters) and three small unassigned populations lacking canonical cell-type markers (Fig. 1B, Fig S2, Table S1).

We first examined two broadly used sentinel markers, *FLG22-INDUCED RECEPTOR-LIKE KINASE 1* (*FRK1*), diagnostic of a transient early response during PTI (Asai *et al*., 2002), and *ISOCHORISMATE SYNTHASE1* (*ICS1*), a hallmark of salicylic acid (SA) synthesis upon pathogen attack (Wildermuth *et al*., 2001). Upon *Pst* DC3000 (EV) challenge, *FRK1* expression peaks at 3 hpi and subsides by 5 hpi, reflecting a defense spike, consistent with previous study using bulk RNA sequencing (Lewis *et al*., 2015). Under *Pst* DC3000 (AvrRpt2) challenge, *ICS1* expression increases steadily until 5 hpi, indicating prolonged SA biosynthesis activation (Fig. 1C).

To generate a set of immune response markers as well as to rule out transcriptomic changes due to protoplasting, we performed bulk RNA sequencing with protoplasts isolated after infection. We identified 897 differentially expressed genes (DEGs) (|log_₂_FC| ≥ 1, FDR < 0.05; 565 up, 332 down; Table S2) from bulk RNA-seq of protoplasts 4 hpi after infection with *Pst* DC3000 (AvrRpt2) when compared with protoplasts from mock samples. Note that comparison of *Pst* DC3000 (EV) to the mock sample yielded only 61 DEGs (Table S2).

We determined an integrated Immunity Score based on the 897 DEGs, by calculating the sum of relative expression of all DEGs. For each cell in our *Pst* DC3000 (AvrRpt2) infected samples, we could thereby map the net transcriptional shift toward activation or repression of immunity (Fig. 1D; Fig. S3). The Immunity Score demonstrated that all cell clusters have the ability to activate an immune response. The Immunity Score was highest in Mesophyll_9 and Mesophyll_8 at 3 hpi, marking these as early response cell clusters.

### Cluster-Modulated Deployment of Defense and Housekeeping Modules under Virulent *Pst* DC3000 (EV) Challenge

To dissect how different types of genes are modulated across different cell populations during virulent challenge, we identified DEGs in each cell cluster (|log_₂_FC| > 0.75, *p adj* < 0.05). In *Pst* DC3000 (EV) infected leaves, most, but not all, DEGs were shared across multiple clusters (Fig. 2A, Table S3-1). Examples for notable genes upregulated specifically after *Pst* DC3000 (EV) infection (Fig. 2B) were the known defense related gene for β-glucosidase 40 (*BGLU40*) (Xu *et al*., 2012; Yamada *et al*., 2020), and AT2G05540, a gene for a glycine-rich protein that had not been linked to defense before (Suh *et al*., 2005). These genes showed distinct temporal and spatial patterns of induction, with *BGLU40* upregulated in the epidermis at 3 hpi and AT2G05540 at 5 hpi.

**Fig. 2.**
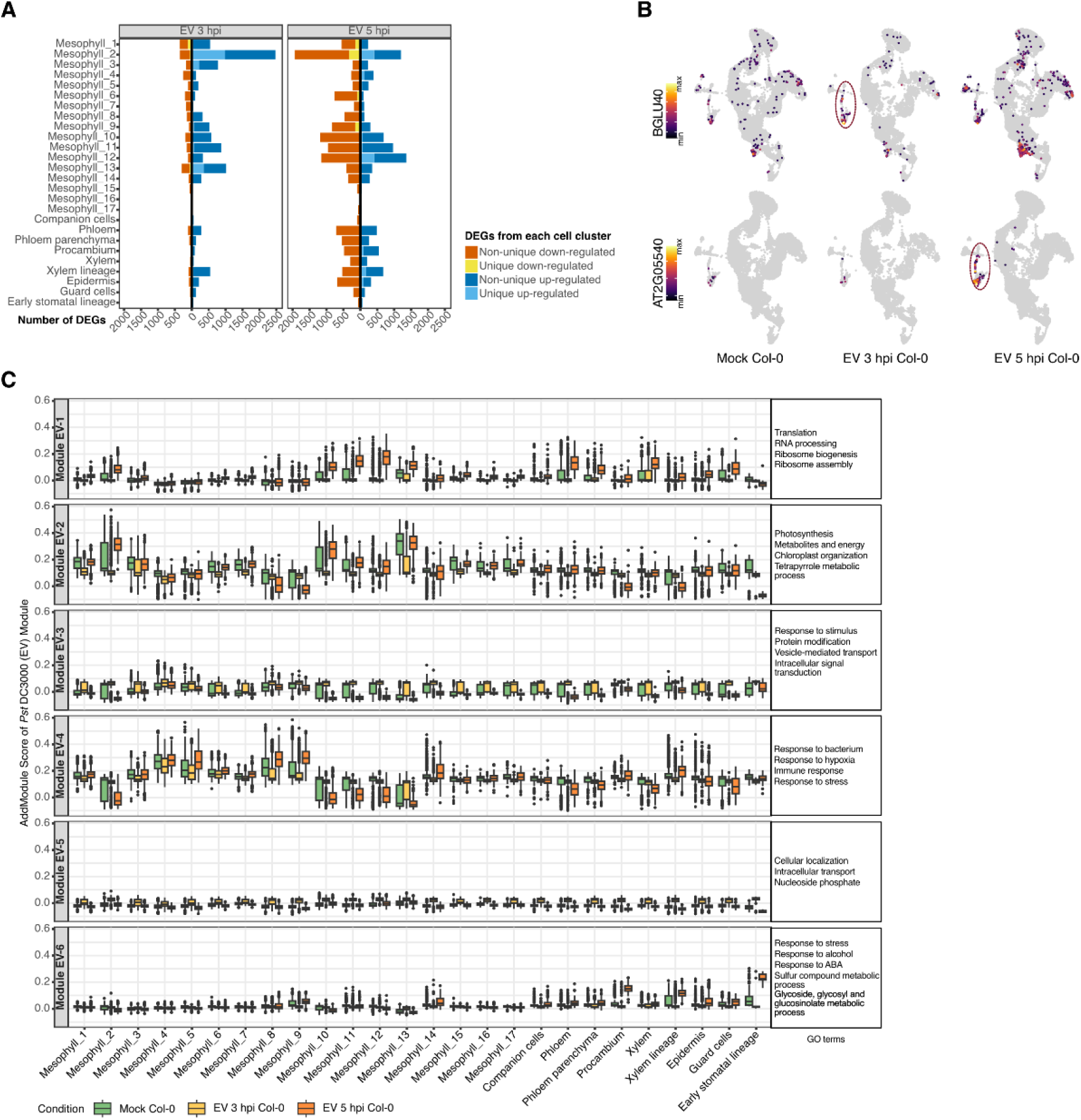
Early dynamic defense response of leaf cell clusters upon *Pst* DC3000 (EV) infection. **A.** Bar plots of DEGs (p-adj < 0.05, |log2FC|>0.75) in each Seurat-defined cell cluster in EV 3 hpi vs. mock (left) and EV 5 hpi vs. mock (right) samples. Bars are stacked by DEG category as indicated on the right. Clusters are ordered by similarity. **B.** UMAP plots of two cluster-specific marker genes, *BGLU40*/AT1G60090 (top row) and AT2G05540 (bottom row), in mock, EV 3 hpi, and EV 5 hpi samples. Color intensity indicates relative normalized expression. Red ovals highlight the clusters in which each gene is most strongly induced upon infection. **C.** Boxplots of module scores for six co-expression modules, calculated with Seurat’s AddModuleScore using DEGs from all cell clusters comparing EV 3 hpi and EV 5 hpi with mock samples (as shown in **A**). The average of module scores per cell cluster is indicated. Color code of samples on the bottom. Top enriched Gene Ontology (GO) terms for each module are listed to the right of each panel. EV is short for *Pst* DC3000 (EV).

To resolve co-regulated functional modules, we performed k-means clustering (Gu *et al*., 2016) on all DEGs identified in *Pst* DC3000 (EV) infected samples (3 hpi vs. mock and 5 hpi vs. mock comparisons for each cell cluster), partitioning genes into six distinct modules (Module EV-1 to EV-6) based on their expression profiles across cell clusters and time points (Fig. 2C; Fig. S4, Table S4-1). Following infection with *Pst* DC3000 (EV), Module EV-2, dominated by photosynthesis and chloroplast-related genes, was transiently down-regulated at 3 hpi before rebounding at 5 hpi in most of the cell clusters, likely reflecting a brief suspension of growth. On the contrary, Modules EV-3 and EV-5, which were enriched for housekeeping Gene Ontology (GO) terms, such as protein modification, vesicle-mediated transport, and nucleoside phosphate, were transiently induced at 3 hpi across most clusters but returned to pre-infection levels by 5 hpi. The defense-related Module EV-4 was downregulated at 3 hpi, but its expression recovered or even became elevated in most mesophyll cells at 5 hpi. In contrast, Module EV-4 is upregulated in Mesophyll_2, _10, _11 and _12 at 3 hpi but becomes downregulated in these clusters at 5 hpi. Finally, Module EV-6, which contains ABA signaling and glycoside/glucosinolate metabolism genes, was exclusively activated at 5 hpi in vascular and early stomatal-lineage cells, pointing to a cell-type specific hormone-mediated stress response (Fig. 2C).

To identify the transcriptional drivers underlying the responses of the different clusters, we inferred gene regulatory networks (GRNs) defined by specific TFs with MINI-EX (Ferrari *et al*., 2022). Regulons controlled by the relevant TFs were ranked by their cell-cluster specificity (borda_clusterRank), and the top ten regulons for each cluster for the mock, *Pst* DC3000 (EV) 3 hpi and *Pst* DC3000 (EV) 5 hpi samples were extracted and then visualized (Fig. S5, Table S5).

There were regulons that were shared across cell types and different infections, but also unique regulons determined by cell type or the response to pathogen infection. In the mock condition, the networks regulated by RHL41/ZAT12, ABR1, MYB74, MYB102, MYB15, and ZAT6 consistently appeared as top-ranked regulons. These six TFs are well-known for their roles in abiotic or biotic-stress responses and growth-related pathways. For example, RHL41 regulates high-light acclimation as well as cold and oxidative stress responses; ABR1, an ABA repressor, interacts with multiple *Pseudomonas syringae* effectors; MYB74 controls osmotic stress to decrease plant growth; and MYB102 increases aphid susceptibility (Iida *et al*., 2000; Zhu *et al*., 2018; Bäumler *et al*., 2019; Schreiber *et al*., 2021; Ortiz-García *et al*., 2022)). Notably, their regulons were specifically enriched in the Mesophyll_2, _10, and _13 clusters, indicating that this set of stress regulators helps define a broader gene regulatory network in uninfected tissue. Upon DC3000 (EV) infection, many of these same regulons remain active, but additional, cell-type specific regulons emerge. For example, MYB96 (Seo *et al*., 2011), a key regulator of wax biosynthesis, defines a top regulon in epidermal clusters at 5 hpi, implicating cuticular reinforcement in epidermal defense.

Taken together, we observed that shared defense and housekeeping gene modules are deployed broadly, but that the onset and intensity of activation of each module are tuned at the cell-cluster level, revealing a gating mechanism by which individual cell populations calibrate their response to pathogen attack. By reconstructing GRNs, we reveal a dual logic whereby common regulons maintain cluster identity while distinct, cell-type-specific networks fine-tune each cluster’s transcriptional response to pathogen challenge.

### Cell Cluster-Dependent Deployment of Defense and Housekeeping Modules under Avirulent *Pst* DC3000 (AvrRpt2) infection

Upon *Pst* DC3000 (AvrRpt2) challenge, the majority of DEGs were shared across clusters, and only a small fraction was truly cluster specific (Fig. 3A, Table S3-2). Among the few DEGs restricted to individual cell clusters, we identified *SCAMP5*, a gene for a membrane trafficking protein that interacts with the TPLATE complex to regulate clathrin-mediated endocytosis and that modulates the localization of aquaporins such as PIP2;1, thereby impacting stress responses including drought tolerance (Yperman *et al*., 2021) (Fig. 5B). The role of SCAMP5 in vesicle trafficking suggests that it may regulate the turnover or localization of defense-related proteins during pathogen infection. Another highly specific DEG was the gene for PIP1;4, an aquaporin that facilitates apoplastic H_2_O_2_ transport during defense against *Pseudomonas (Tian et al., 2016)*. Both *SCAMP5* and *PIP1;4* were specifically activated in guard cells, potentially reflecting a coordinated regulation of membrane trafficking and ROS signaling critical for stomatal defense during early infection stages.

**Fig. 3.**
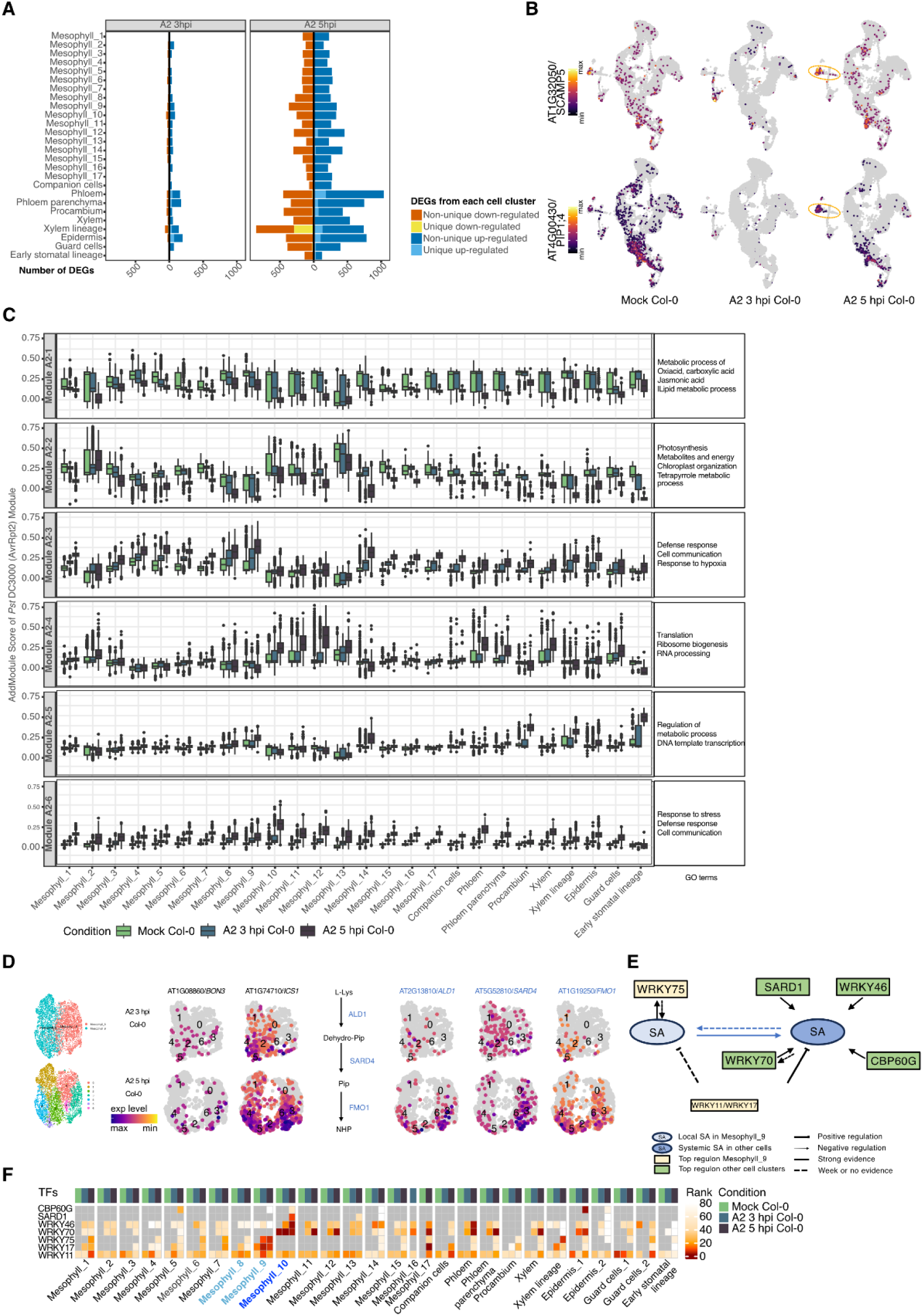
Early responses of leaf cell clusters upon *Pst* DC3000 (AvrRpt2) infection. **A.** Bar plots of DEGs (p-adj < 0.05, |log2FC|>0.75) in each cell cluster in A2 3 hpi vs. mock (left) and A2 5 hpi vs. mock (right) samples. Bars are stacked by DEG category as indicated on the right. **B.** UMAPs plots of two cluster-specific marker genes, *SCAMP5*/AT1G32050 (top row) and *PIP1;4*/AT4G00430 (bottom row) in mock, A2 3 hpi and A2 5 hpi samples. Color denotes relative normalized expression. Red ovals highlight the clusters in which each gene is most strongly induced upon infection. **C.** Boxplots of module scores for six co-expression modules, calculated with Seurat’s AddModuleScore using DEGs from all cell clusters in A2 3 hpi/5 hpi vs. mock samples (as shown in **A**). For each module, the y-axis shows the per-cell score; the x-axis lists all clusters. Color code of samples on the bottom. Top enriched GO terms for each module are listed to the right of each panel. **D.** Subclusters generated from Mesophyll_8 and _9 clusters (left), and the expression of immune-related genes in these subclusters (right). **E.** Diagram of the SA regulatory circuit inferred in Mesophyll_9 versus other clusters. Arrows indicate positive regulation; T-lines indicate repression; dashed arrows indicate weaker or indirect evidence. Key regulons controlled by WRKY75, SARD1, WRKY46, WRKY70, and CBP60G are highlighted, along with feedback between TFs involved in SA biosynthesis and response. **F.** Heatmap of regulon enrichment ranks for seven SA-pathway transcription factors across all clusters and conditions. Each tile’s color (blue→red) indicates the relative rank (1 = strongest enrichment, 80 = weakest) of that TF’s predicted activity in a given cluster in mock (green), A2 3 hpi (light teal), and A2 5 hpi (dark teal) samples. Mesophyll_8, _9 and _10 are annotated in bold. A2 is short for *Pst* DC3000 (AvrRpt2).

To understand the broader functional shifts, we performed k-means clustering on all DEGs, defining gene modules with distinct expression profiles (Fig. 3C and Fig. S6, Table S4-2). Module A2-1, a JA/oxoacid/lipid-metabolism gene set, was progressively repressed between 3 and 5 hpi. Concurrently, the photosynthesis/chloroplast related Module A2-2 was unaffected at 3 hpi but declined by 5 hpi in nearly every cell cluster. In contrast, two defense-related modules, Module A2-3 and Module A2-6, showed robust upregulation across all cell clusters at 3 hpi, with further amplification at 5 hpi, highlighting a broad but intensifying wave of defense.

Building on these module-level dynamics, we identified the cell clusters with the earliest and strongest responses. The integrated Immunity Score, calculated from overall DEG patterns (Fig. 1D), pointed to Mesophyll_9 and _8 as the clusters that respond the earliest to *Pst* DC3000 (AvrRpt2). Further subdivision of these two populations into 7 subclusters revealed specific patterns: the PRIMER cell marker gene *BON3* showed sparse expression in Mesophyll_8 and _9 at 3 hpi and 5 hpi, which may be explained by late induction of *BON3* in PRIMER cells (Nobori *et al*., 2025). *ICS1*, a hallmark of SA biosynthesis, was upregulated at 3 hpi, with expression further concentrating in Mesophyll_8 at 5 hpi (Fig. 3D). Additionally, several key components of systemic acquired resistance (SAR), such as *ALD1*, *SARD4*, and *FMO1*, were induced in the Mesophyll_8 cluster at 5 hpi (Fig. 3D). This localized induction strongly suggests that Mesophyll_8 acts as a "bystander" cell population, actively sensing and amplifying defense signals initiated by adjacent PRIMER cells.

We again used MINI-EX (Ferrari *et al*., 2022) to identify top-ranked regulons in each cell cluster (Fig. 3E and F, Fig. S7, Table S5). Even under pathogen exposure, the key cell identity network, controlled by TF FAMA (AT3G24140) (Ohashi-Ito and Bergmann 2006), remained among the most highly ranked regulon in guard cells.

Focusing on the central defense phytohormone SA, we examined TFs that regulate *ICS1* and *PBS3*, the two key enzymes of SA biosynthesis. The known SA activators *SARD1* and *WRKY46* (Zhang *et al*., 2010; Wang *et al*., 2011) were specifically enriched in Mesophyll_10 together with SA-JA signalling hub TF gene *WRKY70* (Li *et al*., 2004), in *Pst* DC3000 (AvrRpt2) 5 hpi samples (Fig. 3F, Fig. S7).

At the same time, the SA-responsive TF gene *WRKY75*, which promotes SA by antagonizing JA and promoting ROS (Guo et al. 2017, Li et al. 2004), was absent from Mesophyll_9 in the mock treated cells but was dramatically upregulated to define one of the top regulons at both 3 hpi and 5 hpi (Fig. 3E and F). Coupled with the presence of a regulon regulated by GT-3A (AT5G01380) (Fig. S7), a trihelix TF gene previously reported to govern defense in PRIMER cells (Nobori *et al*., 2025), this confirms Mesophyll_9 as a primary, early AvrRpt2-responsive population. Conversely, the negative regulators *WRKY11* and *WRKY17*, which antagonize defense via JA-dependent pathways (Journot-Catalino *et al*., 2006), first appeared in Mesophyll_9, and later also in other clusters, during *Pst* DC3000 (AvrRpt2) infection. The function of negative regulation of SA circuits during early defense remains, however, unclear.

WRKY70 also appeared among the top regulons in Mesophyll_11 and _12, and in epidermal and vascular clusters at both 3 hpi and 5 hpi. A CBP60G related regulon was enriched in the epidermis, consistent with another recent study (Chhillar *et al*., 2025). Together, this suggested that the circuits underlying SA responsiveness and SA biosynthesis are activated in overlapping, but not identical cell populations: in Mesophyll_10, both SA response regulator *WRKY70* as well as the master regulators of SA biosynthesis *SARD1* and *WRKY46* are activated. In contrast, the SA response regulator*WRKY75* is only activated in Mesophyll_9, and the SA biosynthesis regulator *CBP60G* is only activated in the epidermis. Collectively, this set of TFs and the genes they regulate define complex cell-cluster-specific networks that modulate SA biosynthesis, SA response, and SA-JA crosstalk.

Together, these findings reveal a two-tiered regulatory landscape in early plant immunity: core lineage regulons robustly preserve cell-type identity under stress, while concurrently, shared defense and metabolic gene modules are deployed broadly but tuned to different extent in their timing and magnitude by each cell cluster. This demonstrates a sophisticated gating mechanism wherein individual cell populations, through cluster-specific activation of defense-associated transcription factors, calibrate their unique contributions to the overall immune response.

### The Expression Patterns of RLK and NLR Genes Are Tuned by Cell Clusters

To determine in detail whether ETI and PTI components were differently deployed across cell clusters, we examined the expression patterns of key receptors in immunity: RLKs, which include also developmental regulators, and NLRs, which rarely seem to function in processes other than defense. From the known receptor repertoire (Shiu *et al*., 2004), we identified a total of 466 RLK genes, including RLCKs after excluding genes with low expression variance (see Methods), as well as 132 genes encoding Nucleotide-Binding Leucine-Rich Repeat (NLR) proteins as expressed in our data (Table S6). We defined modules for RLKs and NLRs only, as we had done before for all genes.

Mesophyll_4 and _5 had higher basal expression levels of both RLKs and NLRs before pathogen challenge, as demonstrated by their grouping in k-means Module 3 and 4 for RLKs and Module 1 for NLRs (Fig. 4). While the average expression level of RLK genes was elevated at 3 hpi but decreased or returned to mock levels by 5 hpi in the *Pst* DC3000 (EV) samples, a notable exception was in Mesophyll_4, _5, _6, _8, and _9, where the expression of RLK Modules 3 and 4 were distinctly increased at 5 hpi (Fig. 4A and B, Table S6 and S7), indicating a sustained or amplified response capacity in these specific mesophyll subclusters. The RLK Module 4 was upregulated in all cell clusters at 3 hpi, while the RLK Module 1 was primarily induced at 5 hpi in vascular cells. The expression levels of NLRs generally showed a time-dependent increase across many cell clusters upon *Pst* DC3000 (AvrRpt2) infection (Figure 4C and D, Table S6 and S7), suggesting a progressive activation of intracellular defense surveillance.

**Fig. 4.**
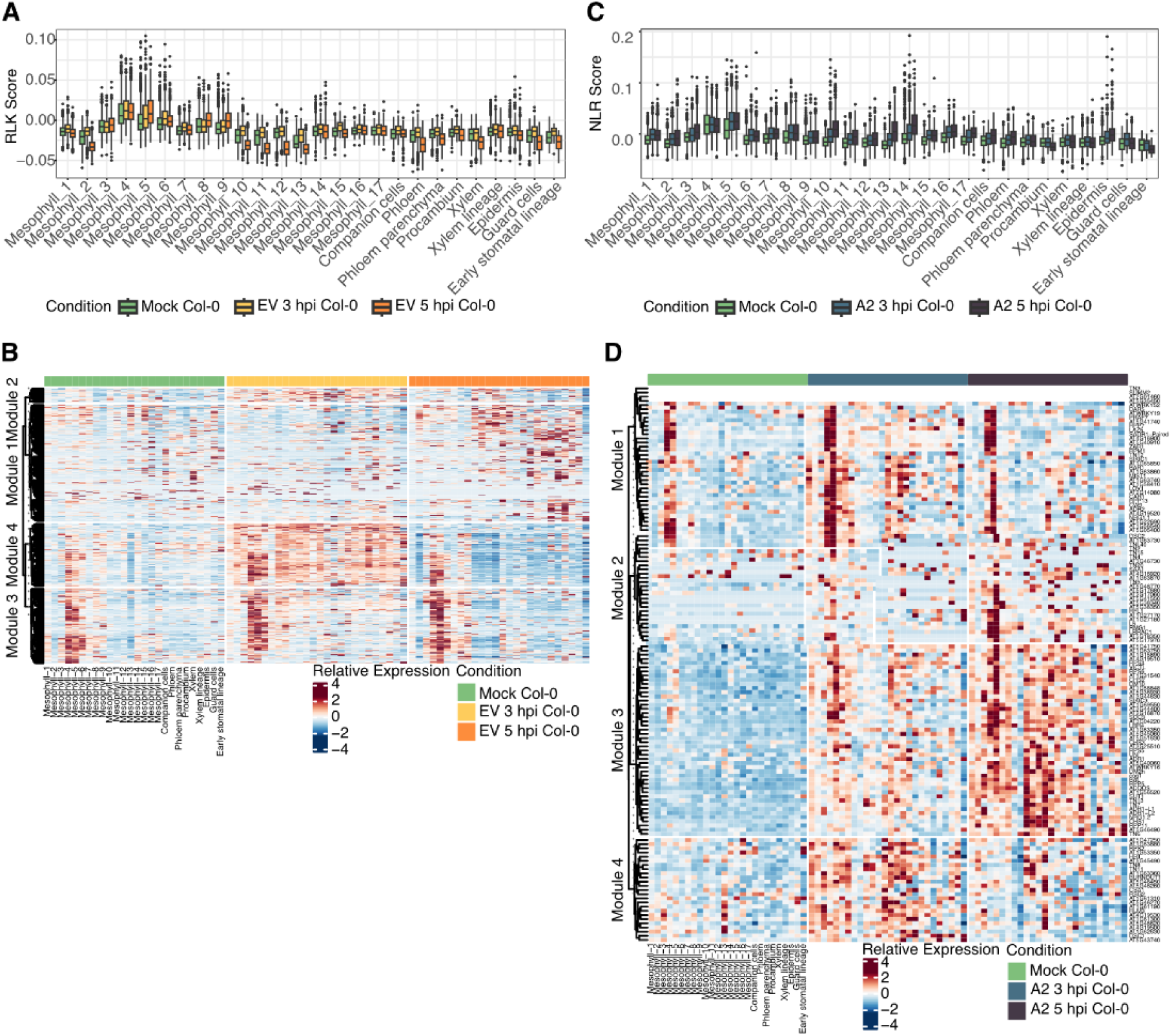
Cluster-resolved expression of RLKs and NLRs. **A.** Line plots showing, for each cell cluster, the average module score of all RLK-encoding genes in mock (green), EV 3 hpi (yellow), and EV 5 hpi (orange) samples. Shaded ribbons represent the standard error of the mean (SEM). **B.** Heatmap of individual RLK genes. Rows are RLKs ordered by hierarchical k-menas clustering, columns are in the same order of cell clusters in mock, EV 3 hpi and EV 5 hpi samples. **C.** Line plots of average module scores for all NLR-encoding genes in each cluster in mock (green), A2 3 hpi (light teal), and A2 5 hpi (dark teal) samples. Shaded ribbons represent the SEM. **D.** Heatmap of individual NLR genes. Rows are NLRs ordered by hierarchical k-means clustering, columns are in the same order of cell clusters in mock, A2 3 hpi and A2 5 hpi samples. EV and A2 are short for *Pst* DC3000 (EV) and *Pst* DC3000 (AvrRpt2), respectively.

Constitutively high expression of immune receptors and defense genes can impose fitness costs, often manifesting as growth retardation or reduced reproductive success (Huot *et al*., 2014; Karasov *et al*., 2017). The uneven expression patterns of RLKs and NLRs may reflect a mechanism that minimizes damage to cell types that are either less important for defense or that are particularly sensitive to activation of defense.

### Spatiotemporal Orchestration of Immune and Growth Programs in Distinct Arabidopsis Leaf Cell Clusters

Returning to all genes, we had found that both *Pst* DC3000 (EV) and *Pst* DC3000 (AvrRpt2) infections induced similar gene modules, including photosynthesis-related modules (Module EV-2 and Module A2-2) and defense-related modules (Module EV-4 and Module A2-3). All these modules had already cell-type-specific expression patterns in mock-treated cells. To systematically characterize the relationship between photosynthesis- and defense-related modules, we defined two core gene sets: a core growth module from the overlap of the photosynthesis-related Module EV-2 and Module A2-2, and a core defense module from the overlap of Module EV-4 and Module A2-3.

Mapping the two core modules onto the different cell clusters, revealed three distinct mesophyll populations: "growth-over-defense" clusters Mesophyll_2, _10, and _13 with high expression of the core growth module and low expression of the core defense module, and "defense-over-growth" clusters Mesophyll_4, _5, _8, and _9, which showed the inverse pattern. A third group of clusters had an intermediate profile (Fig. 5A-B). This observation suggests that mesophyll cells can be specialized in prioritizing either photosynthetic efficiency or immune response.

**Fig. 5.**
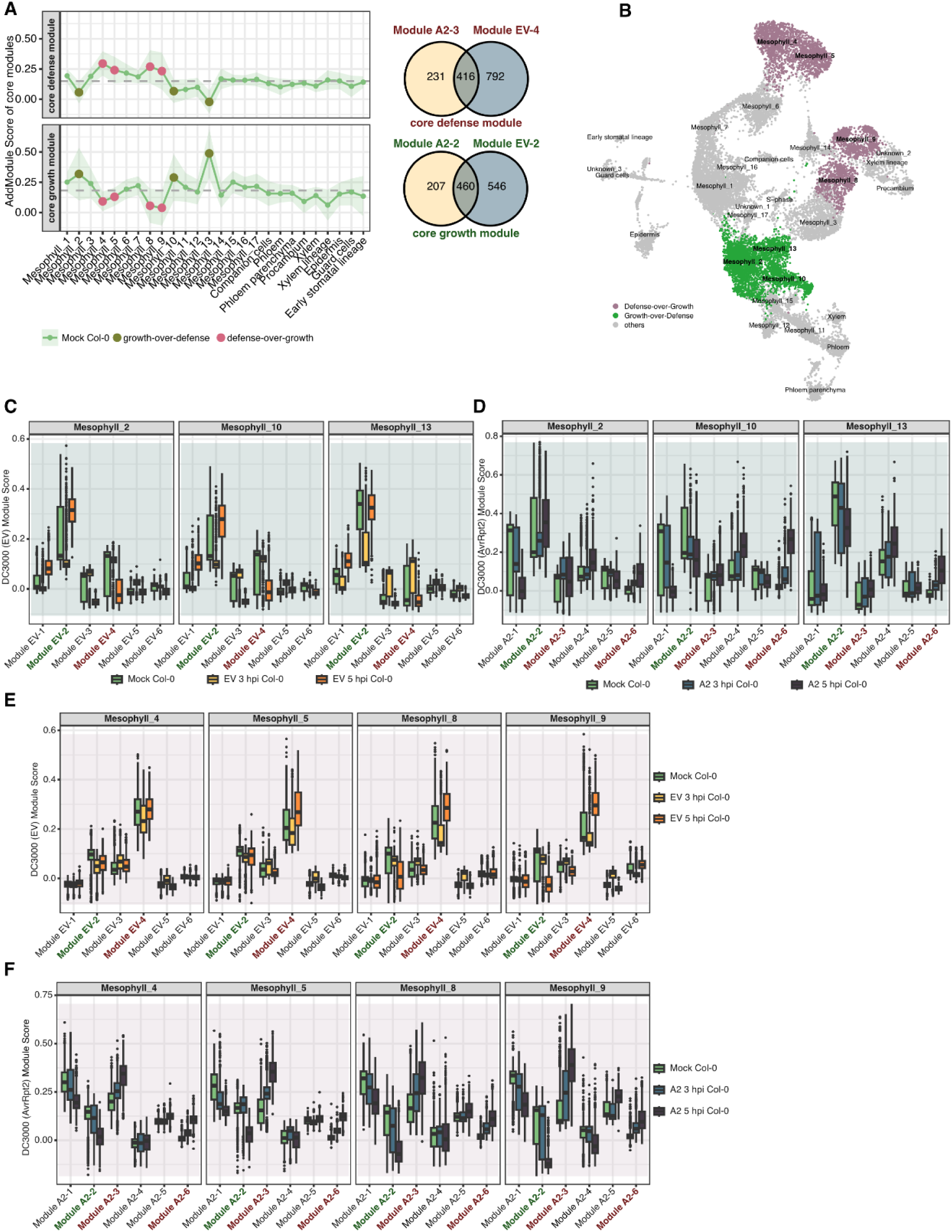
Distinct temporal trajectories during PTI and ETI. **A.** Right: Definition of a core defense module from the overlap of Modules A2-3 and EV-4, and of a core growth module from the overlap of Modules A2-2 and EV-2. Left: Line plots showing the per-cluster average scores for the core defense (top) and core growth (bottom) modules in mock (green) samples. Large colored circles highlight cell clusters with a “defense-over-growth” (magenta) or “growth-over-defense” (olive) phenotype. Shaded ribbons indicate mean ± SEM. **B.** UMAP showing spatial distribution of “growth-over-defense” clusters (green: Mesophyll_2, _10, _13) versus “defense-over-growth” clusters (magenta: Mesophyll_4, _5, _8, _9) in the mock samples; all other clusters in gray. **C-F.** Boxplots of per-cell module scores in the “growth-over-defense” (C, D) and “defense-over-growth” (E, F) cell clusters in (C, E) mock (dark green), EV 3 hpi (gold), EV 5 hpi (orange) and (D, F) *m*ock (dark green), A2 3 hpi (light teal) and A2 5 hpi (dark teal) samples. Within each panel, the growth-related Modules EV-2 and A2-2 are labeled in green, and the defense-related Modules EV-4 and A2-3, A2-6 in red. EV and A2 are short for *Pst* DC3000 (EV) and *Pst* DC3000 (AvrRpt2), respectively.

In “defense-over-growth” cell clusters, defense-related modules were upregulated at 5 hpi upon both *Pst* DC3000 (EV) and *Pst* DC3000 (AvrRpt2) infection (Fig. 5E-F). In contrast, “growth-over-defense” cell clusters showed different behaviors upon *Pst* DC3000 (EV) and *Pst* DC3000 (AvrRpt2) infections. Upon *Pst* DC3000 (EV) infection, these cells only transiently suppressed the growth-related Module EV-2 at 3 hpi (Fig. 5C), and recovered or even upregulated its activity by 5 hpi. The defense-related Module EV-4 was not induced except for modest activation in Mesophyll_13 at 3 hpi and had declined again by 5 hpi (Fig. 5C).

Upon *Pst* DC3000 (AvrRpt2) infection, most cells in both the “growth-over-defense” and “defense-over-growth” clusters activated the defense Modules A2-3 and A2-6 at both 3 hpi and 5 hpi (Fig. 5D and F), and both types of clusters continued to repress the growth-related Module A2-2 also at 5 hpi (Fig. 5E), reflecting a robust ETI-driven shift in gene activity.

These divergent trajectories highlight how the intrinsic heterogeneity of the Arabidopsis leaf transcriptome gates the growth–defense trade-off to determine either early resistance or susceptibility to infection with an avirulent pathogen.

### Potential for Alternative Defense Strategies in the *cue1*-6 Mutant

To investigate how a shift in leaf cell populations affects immunity, we made use of the *cue1*-6 mutant, a reticulate mutant which has fewer mesophyll cells (Streatfield *et al*., 1999; Staehr *et al*., 2014). In our scRNA-seq data, in *cue1*-6 compared to Col-0 wild type, we did notice the proportion of mesophyll cells was lower though did not reach statistical significance (adj *p* = 0.2638) together with a significantly increased proportion of guard cells (adj *p* = 0.0364) (Fig. 6B, Fig. S9).

**Fig. 6.**
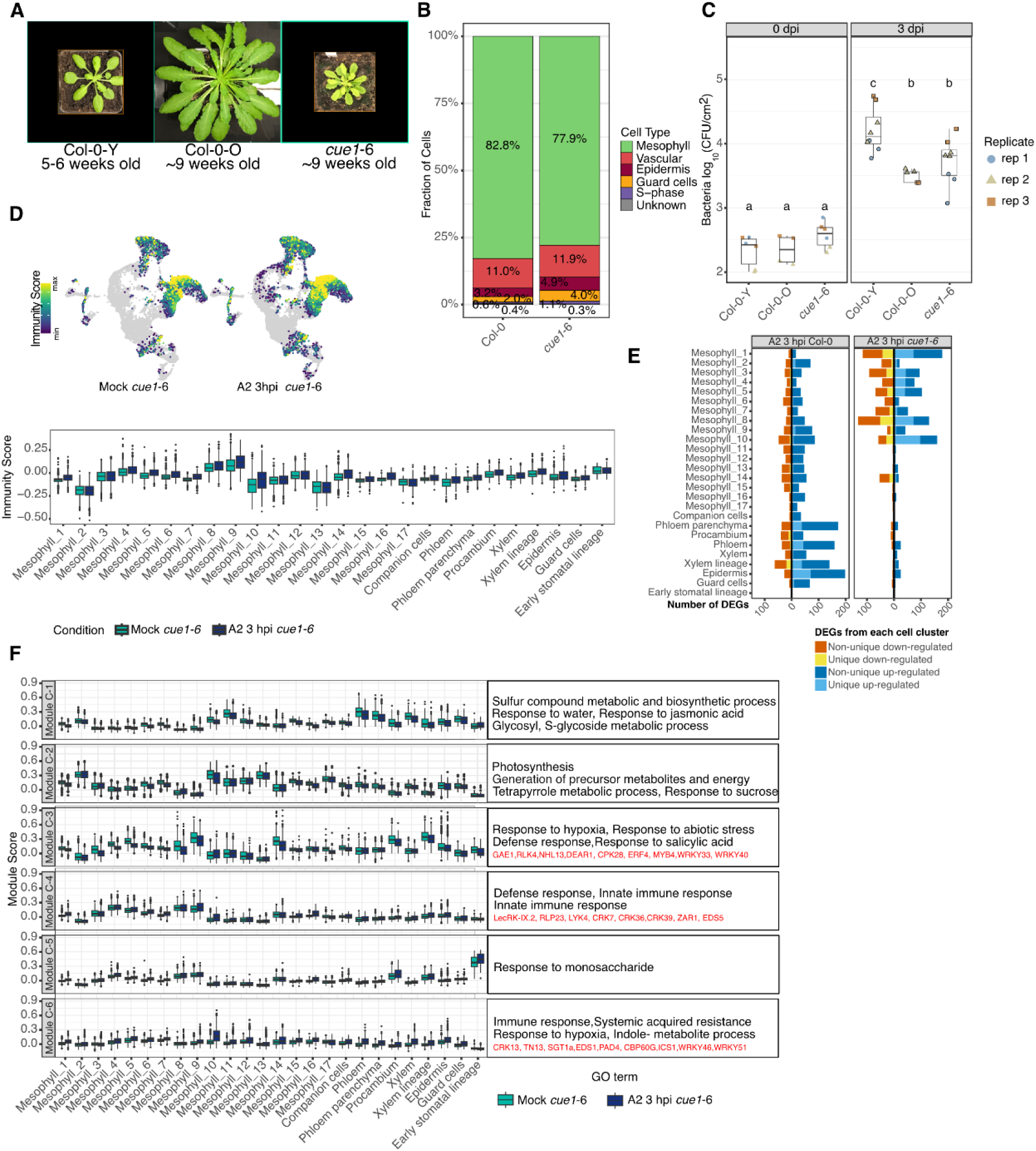
Early ETI responses in *cue1*-6. **A.** Rosette morphology of 5-to-6-week-old Col-0 (Col-0-Y), 9-week-old Col-0 (Col-0-O), and 9-week-old *cue1*-6 plants. **B.** Bar chart of cell-type proportions in Col-0 and *cue1*-6 samples. Colors denote major cell types: mesophyll (green), vasculature (red), epidermis (dark red), guard-cell lineage (orange), S-phase (purple), and unassigned/unknown (gray). **C.** Bacterial growth assay. Leaves were infiltrated with *Pst* DC3000 (AvrRpt2) at OD_600_=0.0001, and bacterial colonization was quantified at 0 or 3 dpi. Each dot represents one individual plant (2 or 3 leaves for Col-0, and 6-8 leaves for *cue1*-6). Colors and shapes represent independent experiment. Log CFU values were compared across sample groups. Different letters indicate significant differences (One-way ANOVA, Tukey’s HSD, p < 0.05). **D.** Immunity Scores highlight the relative expression level of immune-related DEGs (Fig 1D) in each cell (top, UMAP), or in each cell cluster (bottom, boxplot). **E.** Number of DEGs (p-adj < 0.05, |log2FC|>0.75) from each cell cluster of *cue1*-6 plants infected with *Pst* DC3000 (AvrRpt2) at 3 hpi compared with the corresponding cell cluster in mock samples. The left chart shows the number of DEGs in Col-0 after *Pst* DC3000 (AvrRpt2) infiltration at 3 hpi. Bars are stacked by DEG category, as indicated on the right. **F.** Boxplots of average module score values for Modules 1-6 from DEGs in (**E**) in *cue1*-6 mock (green) and *cue1*-6 A2 3 hpi (blue) samples. GO terms and representative top genes for each module are listed alongside the panels. A2 is short for *Pst* DC3000 (AvrRpt2).

Because of their slower growth, we infected *cue1*-6 mutants only after 9 weeks of growth (Fig. 6A). Bacterial growth in *cue1*-6 was comparable to that in older, age-matched Col-0 <u>o</u>ld plants (Col-0-O), but significantly lower than in younger, 5- to 6-week-old Col-0 <u>y</u>oung plants (Col-0-Y) (Fig. 6C), indicating that *cue1*-6 has enhanced resistance relative to wild-type plants of similar size.

The Immunity Score was elevated across nearly all cell clusters in *cue1*-6 infected with *Pst* DC3000 (AvrRpt2) (Fig. 6D). Notably, the number of DEGs unique to specific mesophyll clusters was higher than in Col-0 (Fig. 6E, Table S3-3), suggesting that *cue1*-6 has mechanisms that partially compensate for the altered mesophyll architecture. We categorized DEGs from all cell clusters into six co-expression modules. These modules were similar to modules defined in Col-0 infected with *Pst* DC3000 (AvrRpt2), with Module C-1, enriched for JA response genes, and Module C-2, associated with photosynthesis, being downregulated across most clusters; conversely, Modules C-4 and C-6, enriched for immune response genes, were upregulated in the majority of clusters (Fig. 6F, Fig. S10, Table S4-3). Intriguingly, Modules C-2 and C-5 were enriched for genes involved in sucrose and monosaccharide responses, respectively (Fig. 6F), highlighting potentially an alternative defense strategy. Given that *cue1*-6 has reduced sucrose accumulation (Voll *et al*., 2003), these data suggest that modulation of sugar metabolism and signaling pathways may contribute to the early defense response against pathogens in this mutant.

We also examined the expression patterns of critical TF genes involved in SA and JA signaling across different cell clusters. Key SA-related WRKY regulons, including those defined by WRKY75, WRKY17, WRKY11, WRKY70, and WRKY46, were detected in various *cue1*-6 cell clusters, with expression patterns similar to those observed in Col-0 wild type. For example, the WRKY46 regulon showed specific induction in Mesophyll_10, while WRKY75 and WRKY17 regulons were predominantly activated in Mesophyll_9 after infection (Fig. S11, Table S5). Additionally, the PRIMER cell marker *GT-3A* (Nobori *et al*., 2025) related regulon, was enriched in Mesophyll_4, _8, and _9 in both mock and *Pst* DC3000 (AvrRpt2) infected samples, similar to its distribution in Col-0 wild type (Fig. S11, Fig S7).

Regarding JA signaling, the regulon controlled by the master TF MYC2, a well-established activator of JA-mediated defense responses, was among the top 10 regulons primarily induced in phloem tissues, but it was notably inactive in Mesophyll_14 and epidermal clusters (Fig. S11). Conversely, the regulon controlled by JAM2, a negative regulator of JA signaling that antagonizes MYC2 activity (Sasaki-Sekimoto *et al*., 2013), was induced in the Mesophyll_4, _5 and _16 clusters. This complementary and spatially distinct expression patterns of MYC2 and JAM2 regulons suggests a complex, possibly non-cell-autonomous antagonistic regulation of JA signaling across cell types.

Together, these results demonstrate that despite developmental constraints, *cue1*-6 mutants are resilient in their immune responses. Most core immune modules are similar to those in wild type, but we also see a potentially alternative strategy, altered sucrose signaling. The cell cluster-specific distribution of these TF regulons underscores the importance of cell-type context in hormone-mediated defense.

## Discussion

A perennial topic in plant immunity is the balance between growth and defense. Clearly, different cell layers, such as epidermis and mesophyll, play very different roles in defense, and the growth-defense trade-off therefore is likely to play out differently in these layers (Wyrsch *et al*., 2015). In addition, there is division of labor among different mesophyll cells when it comes to processes such as photosynthesis (Xia *et al*., 2022). Here, we tested the hypothesis that growth-defense tradeoffs are managed in a cell-type-specific manner.

### Intrinsic Mesophyll Heterogeneity Driving Growth-Defense Trade-offs and Shaping Infection Outcomes

The major cell type of leaves, mesophyll cells, features heterogeneous transcriptomic profiles influenced by physical location of cells and their developmental state (Xia *et al*., 2022; Guo *et al*., 2025). Previous single-cell transcriptome studies have already described cell-type specific heterogeneity of pathogen responses in the Arabidopsis leaf (Zhu *et al*., 2023; Delannoy *et al*., 2023; Tang *et al*., 2023; Nobori *et al*., 2025; Chhillar *et al*., 2025). Our study, which specifically focused on mesophyll cells, identified resilient cell populations that quickly reinitiate programs supporting growth after pathogen attack.

We found that specific cell clusters already expressing defense-related genes, including key cell surface receptors and intracellular receptors (Fig. 4), mount stronger and earlier immune responses upon infection with the avirulent pathogen *Pst* DC3000 (AvrRpt2). Conversely, growth-over-defense cells are delayed in the activation of defense (Fig. 5D and F). These observations suggest that intrinsic transcriptional states dictate the speed and magnitude of immune responses in different cell types.

Our data also illuminate how this trade-off is expressed in the face of different challenges. Compared to the virulent *Pst* DC3000 (EV) strain, the avirulent *Pst* DC3000 (AvrRpt2) strain both suppresses photosynthetic activity and activates immunity more uniformly across mesophyll clusters, while the virulent *Pst* DC3000 (EV) strain leads to a more nuanced response. The expression of photosynthesis-related genes is derepressed in growth-over-defense cells by 5 hpi, while defense gene expression is prioritized in defense-over-growth cells (Fig. 5). This observation suggests that virulent pathogen strategies apparently involve selective manipulation of host cellular priorities, an insight with potentially profound implications for engineering durable plant resistance that minimizes growth penalties.

The origins of this intrinsic heterogeneity, whether predominantly developmental, metabolic, or epigenetic, remain an open question worthy of future investigation. A priority should be the identification of TFs and the GRNs they regulate or other signaling factors that connect growth and defense.

### Spatiotemporal Coordination of Immune Modules Determines Resistance

Optimal resistance requires precise temporal and spatial orchestration of core defense modules. While Zhu and colleagues (2023) identified distinct immune and susceptible mesophyll states at later infection stages (16-48 hpi), our analysis reveals dynamic transcriptional reprogramming during the early phases of infection (3-5 hpi). *Pst* DC3000 (AvrRpt2) induces progressive amplification of defense signals, whereas the virulent *Pst* DC3000 (EV) strain is able to dampen initial responses at later time points (Fig. 2 and 3). Notably, photosynthesis genes are rapidly suppressed in resistant interactions, with the proviso that we do not know whether mesophyll cells recover metabolic activity once defense has subsided (Fig. 5).

We identified the Mesophyll_9 cluster as a putative PRIMER cell population and Mesopyll_8 as bystander cells (Nobori *et al*., 2025). Apparent PRIMER cells in our data set are characterized by expression of the TFs GT-3A as well as WRKY75 and WRKY11/17, well-known SA regulators (Fig. 3). PRIMER cells initiate localized cell death while bystander cells express systemic immunity components (Nobori *et al*., 2025). Given that SA promotes cell survival (Zavaliev *et al*., 2020), while PRIMER cells are destined to execute cell death, the enrichment of the SA-response repressors WRKY11/17 in the Mesophyll_9 cluster and their absence in the SA-producing Mesophyll_10 cluster suggests tight spatial control of the execution of cell death (Fig. 3). Future spatial transcriptomics experiments could elucidate how these populations physically interact, and how their precise spatial arrangement dictates immunity outcomes.

### Induced and Natural Genetic Variation as Untapped Resources for the Study of Plant Immunity

Natural and induced genetic variants of plants have been essential for dissecting the mechanisms underlying plant immunity, yet their potential for understanding immunity at single-cell resolution remains largely untapped. Bulk transcriptomic studies in Arabidopsis have demonstrated how phytohormone mutants can be used to reveal layers of immune responses: the *dde2 ein2 pad4 sid2* quadruple mutant, impaired in JA/ethylene/PAD4/SA signaling, has informed how we think about gene regulation during PTI and ETI (Mine *et al*., 2018). Analyses of helper NLR mutants have revealed redundant as well as unequal contributions of these factors to PTI- and ETI-induced transcriptional changes (Saile *et al*., 2020).

Genetics has already been used on the pathogen side for single-cell analyses, for example in a recent study that contrasted virulent and avirulent pathogens and that also compared two distinct effectors, AvrRpt2 and AvrRpm1, (Nobori *et al*., 2025). This study, as well as our study, have revealed genes that appear to control the behavior of different cell types and clusters, and scRNA-seq analysis of plants mutant for these regulators will likely be very informative. In addition, while bulk RNA-seq experiments have shown broadly similar patterns of early transcriptional reprogramming induced by different PAMPs or bacterial strains (Bjornson *et al*., 2021; Maier *et al*., 2021; Keppler *et al*., 2025), only single-cell approaches can reveal how individual cell type decodes these signals. Another avenue will be the comparison of CNL- and TNL-triggered defenses.

We have studied here the *cue1* mutant, which has altered leaf architecture and which is impaired in the shikimate pathway (Li *et al*., 1995; Voll *et al*., 2003). We found *cue1* to mount a robust ETI across all mesophyll clusters, but also, unexpectedly, to engage a sucrose module in the early hours of infection (Fig. 6D and F). These results highlight that developmental status and metabolic context can profoundly shape the immune landscape, potentially activating alternative defense strategies based on the plant’s physiological condition. This bodes well for the use of natural genetic variation to better understand plant immunity at the single-cell level, given what is already known from bulk RNA-seq experiments (Corwin *et al*., 2016).

Looking forward, an important question will be how robust core PTI and ETI GRNs are in different mutant backgrounds or different environments. By systematically profiling mutants and natural accessions with altered environmental responses, growth, metabolism or development, we can map how these processes are integrated with plant immunity, potentially uncovering previously underappreciated points of cross-talk. We believe that single-cell driven knowledge will enhance the field’s ability to engineer crops with better balance between optimal yield and pathogen resistance.

## Material and Methods

### Reagents and Kits

The following reagents and kits were used: Chromium Next GEM Single Cell 3’ Kit v3.1 (10x Genomics,PN-1000268), Chromium Next GEM Chip G Single Cell Kit (10x Genomics, PN-1000120), TWEEN 20 (Sigma-Aldrich, P1379), SPRIselect Bead-Based Reagent (Beckman Coulter, B23317), Low TE Buffer (10 mM Tris-HCl pH 8.0, 0.1 mM EDTA) (Thermo Fisher Scientific, J75793.AP), glycerol (Sigma-Aldrich,G5516), RNeasy Plant Mini Kit (Qiagen, 74904), diethiothreitol (DTT) (Thermo Fisher Scientific, R0861), DNase I, RNase-free (Thermo Fisher Scientific, EN0521), diethyl pyrocarbonate (DEPC) (Sigma-Aldrich, D5758), RNase inhibitor/RNaseOUT (Invitrogen,10777019), MES hydrate (Sigma, 69890), D-Mannitol (Sigma, M1902), Calcium chloride (ROTH, A119.1), potassium chloride (ROTH, 6781.1), magnesium chloride hexahydrate (ROTH, 2189.1), sodium chloride (ROTH, 3957.1), Cellulase-RS (Duchefa Biochemie, C8003) and Macerozyme-R10 (Duchefa Biochemie, M8002).

### Plant Material and Growth Conditions

Seeds of the Arabidopsis reference accession Col-0 and the *cue1*-6 mutant in the same background (kindly provided by Dr. R. E. Häusler, University of Cologne) were surface-sterilized with 30% commercial bleach and stratified at 4°C for one week before sowing on soil. Seven- to ten-day-old seedlings were transplanted into individual pots and grown at 23°C, 8 h light /16 h dark. Unless otherwise noted, Col-0 plants were five to six weeks old and *cue1*-6 plants nine weeks.

### Bacterial Infections and Bacterial Growth Measurements

*Pseudomonas syringae* pv. tomato (*Pst*) DC3000 (empty vector, EV) and *Pst* DC3000 (AvrRpt2) were cultured overnight at 28°C in LB medium with kanamycin (50 mg/L). Cells were pelleted by centrifugation, washed, and resuspended in 10 mM MgCl_₂_. For RNA-seq samples, suspensions were adjusted to OD_₆₀₀_=0.005; for bacterial growth curves, to OD_₆₀₀_=0.0001 to minimize hypersensitive cell death. The abaxial surfaces of leaves were infiltrated with a needle-less syringe, blotted dry, and harvested at 3 hpi and 5 hpi for scRNA-seq samples, at 4 hpi for bulk RNA-seq samples, or at 0 dpi and 3 dpi for bacterial growth measurements. Each treatment comprised three biological replicates (2-3 leaves per Col-0 plant and 3-6 leaves per *cue1*-6 plant).

For measurements of bacterial growth, leaf discs were harvested with a 6-mm-diameter punch on day 0 (immediately after infiltration) and day 3. For day 0 samples, six leaves from two plants were pooled (two discs per leaf), and four discs were combined as one sample. For day 3, four discs from one plant (two discs per leaf) constituted a sample. Discs were ground in 200 µl infiltration buffer, serially diluted (10×, 100×, 1000×), and 10 µl of each dilution was spotted on LB agar plates containing kanamycin (50 µg/ml). Plates were incubated at 28°C for 48-72 h, and bacterial growth was quantified as colony-forming units per leaf area (CFU/cm²).

### Protoplast Isolation

Protoplasts were isolated as described, with sucrose centrifugation to remove debris (Yoo *et al*., 2007; Jeong *et al*., 2021). Leaves were incubated in digestion buffer (1.5% Cellulase-RS, 0.3% Macerozyme R10, 20 mM MES pH 5.7, 0.6 M mannitol, 20 mM KCl, 10 mM CaCl_₂_, 1% BSA) for 1 h at room temperature with gentle agitation. After low-speed centrifugation (150 × *g*, 5 min, 4°C), the pellet was resuspended in MMC solution (10 mM MES, 0.47 M mannitol, 10 mM CaCl_₂_) and layered over 6 mL of 6 M sucrose. Following centrifugation (80 × *g*, 9 min), the middle layer was recovered, washed in 0.5 M mannitol, and pelleted (150 × *g*, 5 min, 4°C). Protoplasts were resuspended in a solution with 0.25 M mannitol and 3 mM sucrose for scRNA-seq. For bulk RNA-seq, cells were collected prior to sucrose gradient centrifugation.

### Bulk RNA-seq and Analysis

Total RNA was extracted from frozen protoplast pellets using the RNeasy Plant Mini Kit (Qiagen). Libraries were prepared from 500 ng RNA as described (Cambiagno *et al*., 2021) and sequenced on an Illumina NextSeq 2000 instrument with 100 bp single-end reads. Reads were aligned to the TAIR10 genome (Araport11 GTF) (Cheng *et al*., 2017) using the nf-core/rna-seq pipeline (v2.0.0) with default parameters and STAR-RSEM for alignment and quantification. Gene-level counts were processed in DESeq2 v1.46.0 (Love *et al*., 2014). Genes with total counts ≤5 were removed, data were rlog-normalized, and DEGs were identified with a Wald test (FDR < 0.05, |log_₂_ FC| ≥ 1). P-values were adjusted for multiple testing using the Benjamini-Hochberg procedure.

### ScRNA-seq and Processing

Protoplasts were counted by light microscope and loaded onto the 10× Chromium Controller using Next GEM 3′ v3.1 reagents. Libraries were sequenced on an NextSeq 2000 instrument (90 bp read 2, dual index). Reads were processed with the nf-core/scrnaseq pipeline v2.0.0 (Cell Ranger v7.0.0) against the Arabidopsis TAIR10 reference.

Raw UMI matrices were imported into Seurat v4 (Hao *et al*., 2021). Only cells with nCount_RNA >1000, 500< nFeature_RNA <8000 and chloroplast transcripts <15% were retained. To identify and remove putative doublets, we used DoubletFinder v2.0.3 (McGinnis *et al*., 2019).

### ScRNA-seq Data Integration and Clustering

Raw counts were first normalized and variance-stabilized using Seurat’s SCTransform (Hafemeister & Satija, 2019). To integrate across batches and conditions, we used Seurat’s anchor-based reciprocal PCA (RPCA) workflow, followed by IntegrateData to produce an “integrated” assay. Clustering analysis was performed with RunPCA (npcs=50), followed by FindNeighbors (). Cell clusters were identified with FindClusters () at a resolution of 0.8.

Uniform manifold approximation and projection (UMAP) visualization was generated from the Harmony embeddings (reduction=’harmony’, dims=1:30). We ran RunUMAP() once on the Harmony embeddings and used the resulting UMAP coordinates for all figures. The resulting Seurat object (seurat.obj including umap_1 and umap_2) is provided in the Supplementary Data.

### Identification of DEGs in scRNA-seq Data and Gene Module Clustering

For each cell cluster, DEGs between mock and infection treatment of the same genotype (Col-0 or *cue1*-6) were identified with FindMarkers on SCT assay. All DEGs (adj p < 0.05, |log_₂_ FC| > 0.75) were combined and partitioned into six co-expression modules by k-means clustering (ComplexHeatmap) (Gu *et al*., 2016), using Euclidean distance and k=6 for Figure 2 and Figure 3, and k=4 for Figure 4. For the heatmaps of RLK and NLR gene expression, gene-level variance was calculated across all samples, and the bottom decile (including genes with zero or near-zero expression in all conditions) was excluded. This conservative threshold was used to preserve condition-specific signals while reducing noise.

### Gene Regulatory Network Inference

MINI-EX (Ferrari *et al*., 2022) was run as described (Cao *et al*., 2023) on individual Seurat objects and corresponding DEG lists for each cell cluster. Regulons were ranked by “borda_clusterRank” specificity, and the top 10 regulons in each cell cluster were visualized with presence/absence heatmaps. Note that there are two cell clusters each for epidermis and guard cells (epidermis_1 and _2, guard cell_1 and _2).

### Statistical Analysis of RLK, NLR Expression Scores and Immunity Score

To assess changes in RLK expression scores in DC3000 (EV) infected samples, NLR expression scores and the Immunity Score in Pst DC3000 (AvrRpt2) infected samples across treatments within each cell cluster, we performed pairwise comparisons between treatments using the Wilcoxon rank-sum test. Analyses were conducted independently for each cell cluster using the rstatix R package (v 0.7.2). To account for multiple hypothesis testing, p-values were adjusted using the Benjamini-Hochberg (BH) method to control the False Discovery Rate (FDR). A p.adj value <0.05 was considered statistically significant.

## Data Availability

Sequencing reads have been deposited in the European Nucleotide Archive (ENA) (PRJEB94524 for scRNA-seq, PRJEB94525 for bulk RNA-seq). Scripts, code and the scRNA-seq processed dataset are available at 10.5281/zenodo.16533111.

## Acknowledgements

We thank Dr. R. E. Häusler (University of Cologne) for providing *cue1*-6 mutant seeds. We thank Christian Hartman for help with plant growth, Tom Denyer from the Timmermans lab for valuable suggestions for scRNA-seq analyses, and all members of the Weigel lab for discussion. Supported by DFG SFB 1101 (M.T. and D.W.), DFG TRR356 (D.W.), the Novo Nordisk Foundation (Novozymes Prize, D.W.) and the Max Planck Society (D.W.).

## Author contributions

S.W. and D.W. planned and designed the research. S.W. and H.G. performed experiments. S.W., I.B., and P-J.W. conducted data analysis. S.W. and D.W. wrote the manuscript. S.W., I.B., P-J.W, H.G., M.T. and D.W. reviewed and edited the manuscript and provided discussions. S.W. and D.W. are corresponding authors.

## Supplementary Material

### Supplementary Figures

**Fig. S1.**
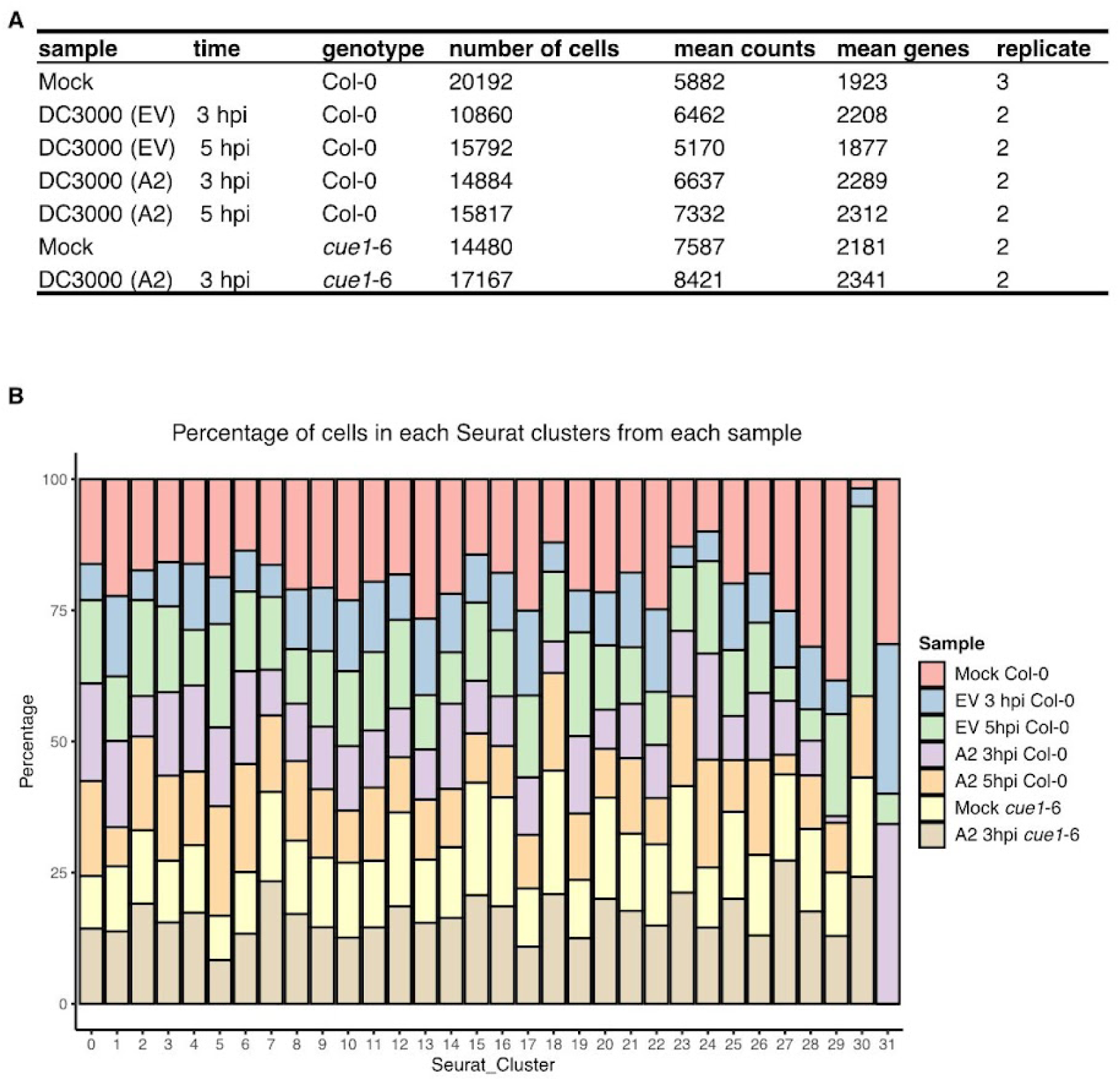
Overview of the scRNA-seq experimental design. **A.** Summary of scRNA-seq data from samples at 3 and 5 hours post infiltration (hpi) of mock, *Pst* DC3000 (EV) and *Pst* DC3000 (AvrRpt2) in Col-0 wild-type and *cue1*-6 mutant plants. Mean counts are mean counts of unique molecular identifiers (UMIs) per cell, mean genes are mean number of genes for which transcripts were detected per cell. **B. Cluster composition by sample.**The proportion of cells from samples described in **A** in each cluster. EV and A2 are short for *Pst* DC3000 (EV) and *Pst* DC3000 (AvrRpt2), respectively.

**Fig. S2.**
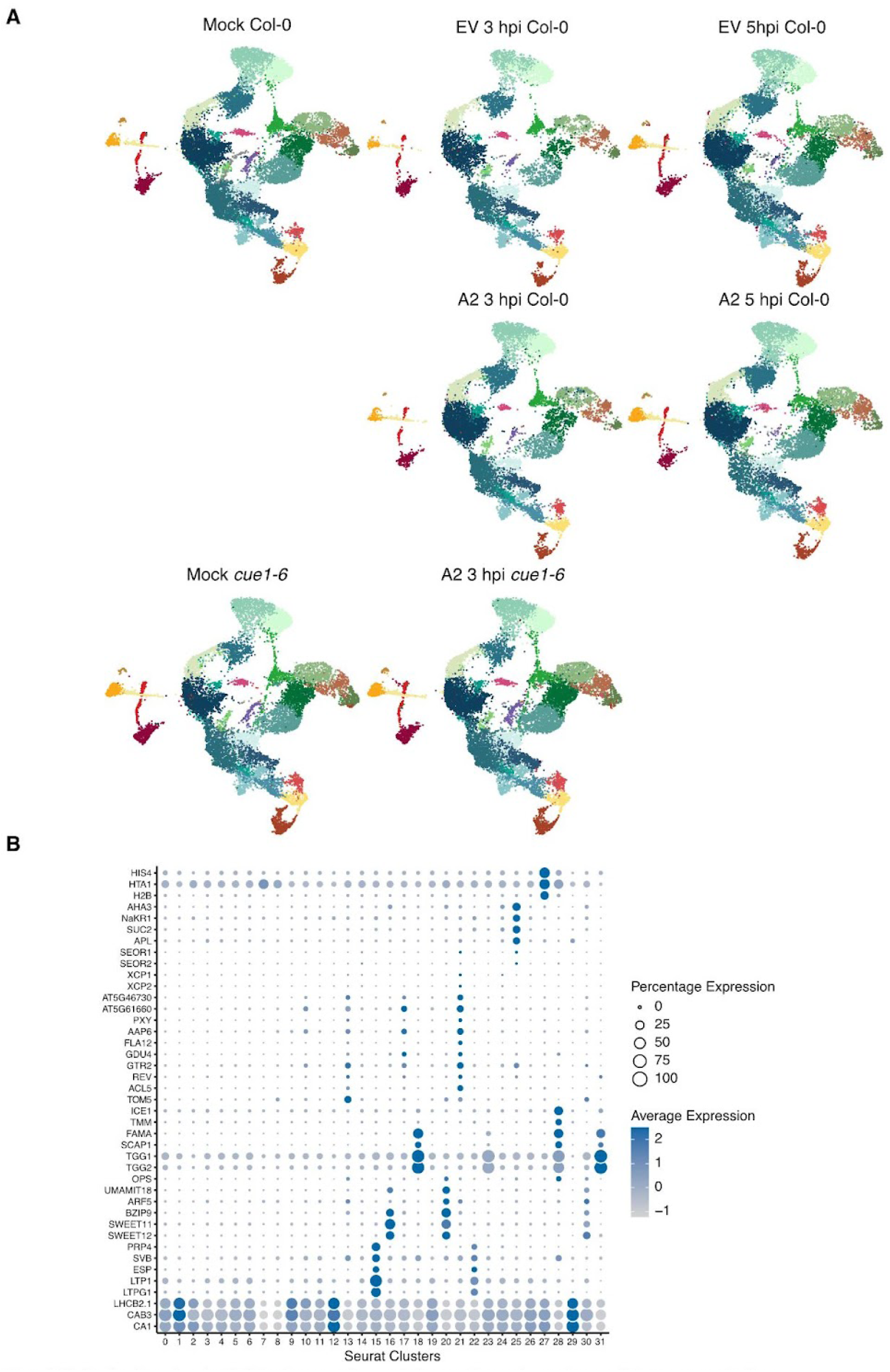
Single-cell atlas of individual samples and marker genes for cell-type annotation. **A.** Uniform Manifold Approximation and Projection (UMAPs) for each sample, with biological replicates combined. EV and A2 are short for *Pst* DC3000 (EV) and *Pst* DC3000 (AvrRpt2), respectively. **B.** Dot plot of known cell type marker genes across Seurat clusters: dot size indicates the fraction of cells expressing each gene, and color represents scaled average expression. Table S1 shows the marker genes and corresponding cell type.

**Fig. S3.**
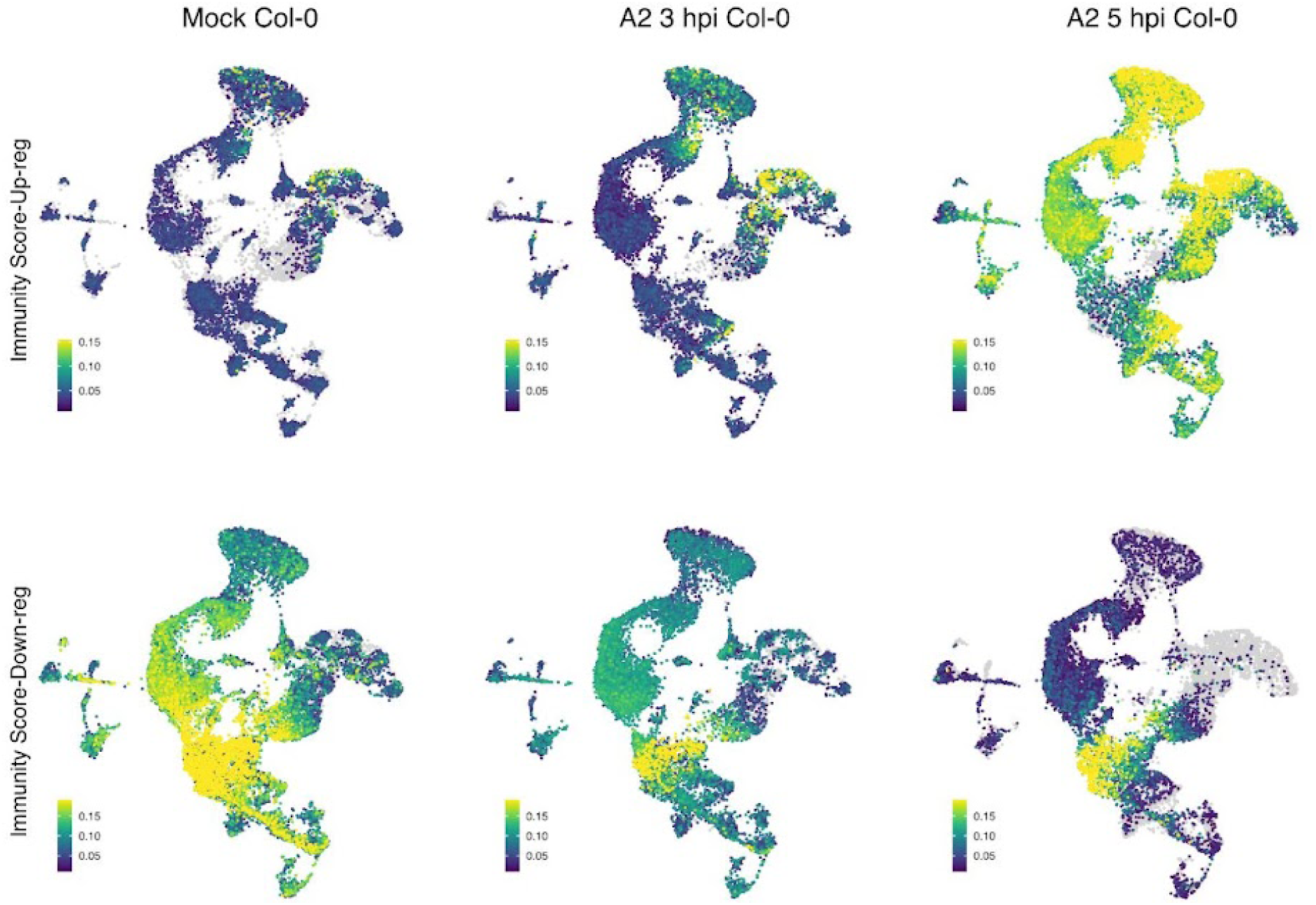
Single-cell distribution of defense-response signatures. UMAPs colored by an Immunity Score calculated for each cell as the mean scaled expression of immune-related DEGs identified by bulk-RNA-seq (see Fig. 1). **Top row:** score based on genes up-regulated by AvrRpt2 (*Up-DEG score*). **Bottom row:** score based on genes down-regulated by AvrRpt2 (*Down-DEG score*). Yellow denotes cells with higher average expression of the indicated gene set, and dark blue denotes lower expression.

**Fig. S4.**
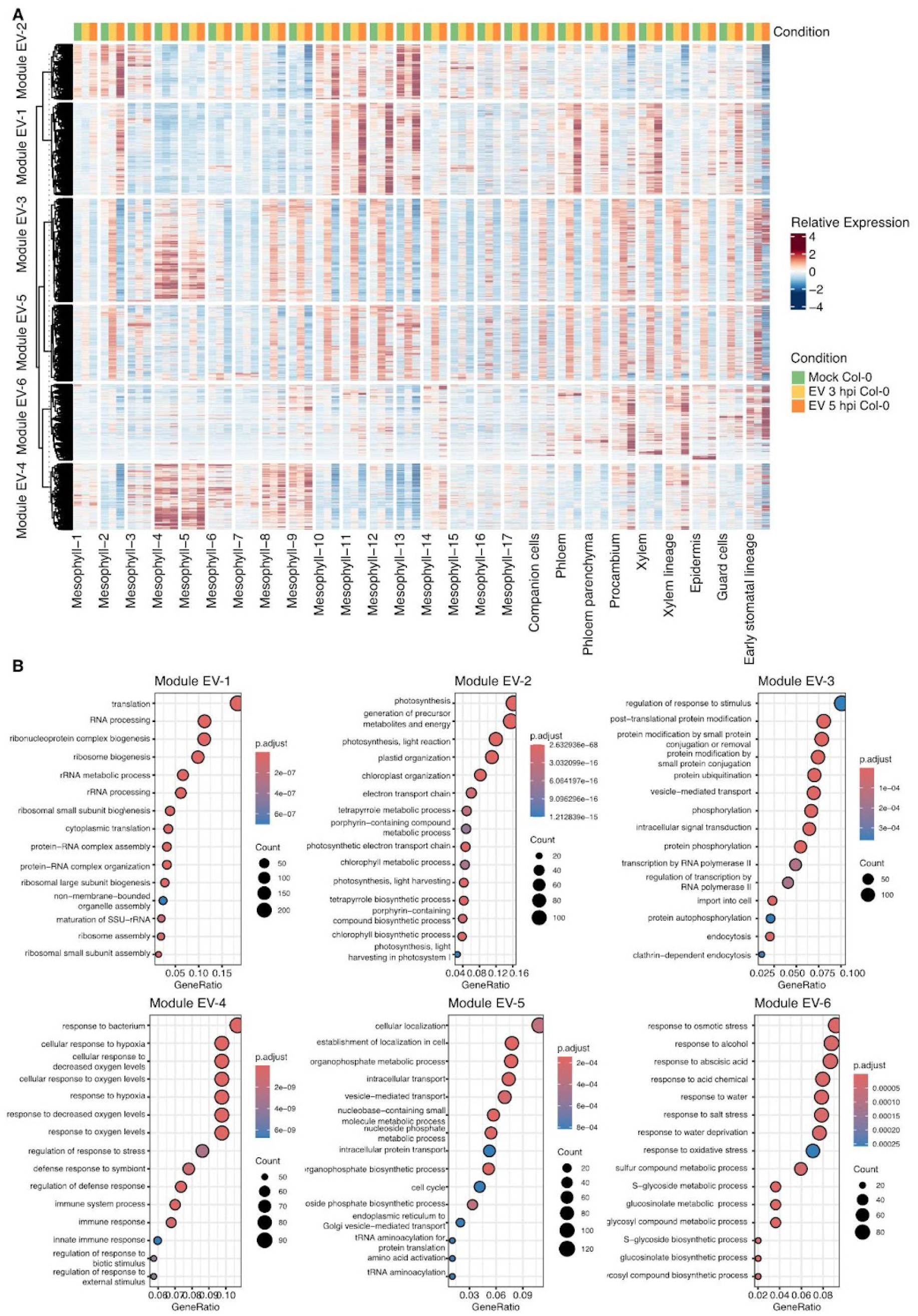
Gene modules in Pst DC3000 (EV) infected Col-0 samples. **A.** Heatmap of DEGs from all cell clusters of EV 3 hpi/5 hpi vs. mock comparisons. DEGs were k-mean clustered into six co-expression modules (Modules EV-1 to EV-6) based on their expression pattern. Columns correspond to the cell clusters; the colored rectangles on top indicate sample types, as indicated on the right, mock (green), EV 3 hpi (yellow) and EV 5 hpi (orange). Heatmap colour shows the relative expression values (red = high, blue = low), as indicated on the right. **B.** Functional enrichment of each module. Dotplots show the top enriched GO terms per module (ClusterProfiler, Benjamini–Hochberg FDR). Dot size represents the number of module genes in the term, and color encodes the adjusted *P* value. EV is short for Pst DC3000 (EV).

**Fig. S5.**
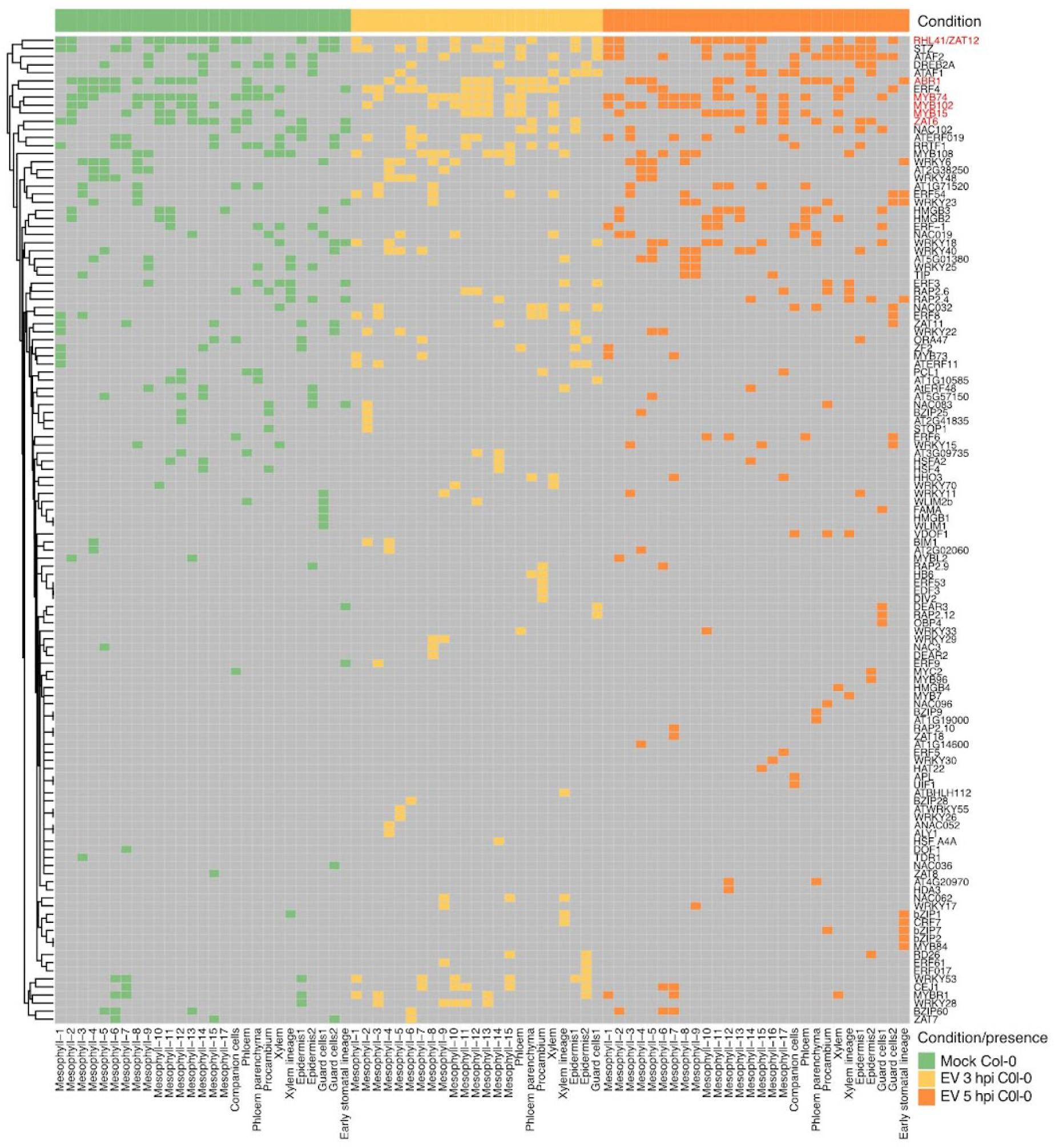
Distribution of the top 10 regulons across cell clusters in Pst DC3000 (EV) infected Col-0 samples. Presence/absence heatmap summarising the ten highest-ranking specific regulons detected in each cell cluster. The color bar above indicates the sample: mock (green), EV 3 hpi (yellow) and EV 5 hpi (orange). Red rectangles highlight the TFs related to stress and growth.

**Fig. S6.**
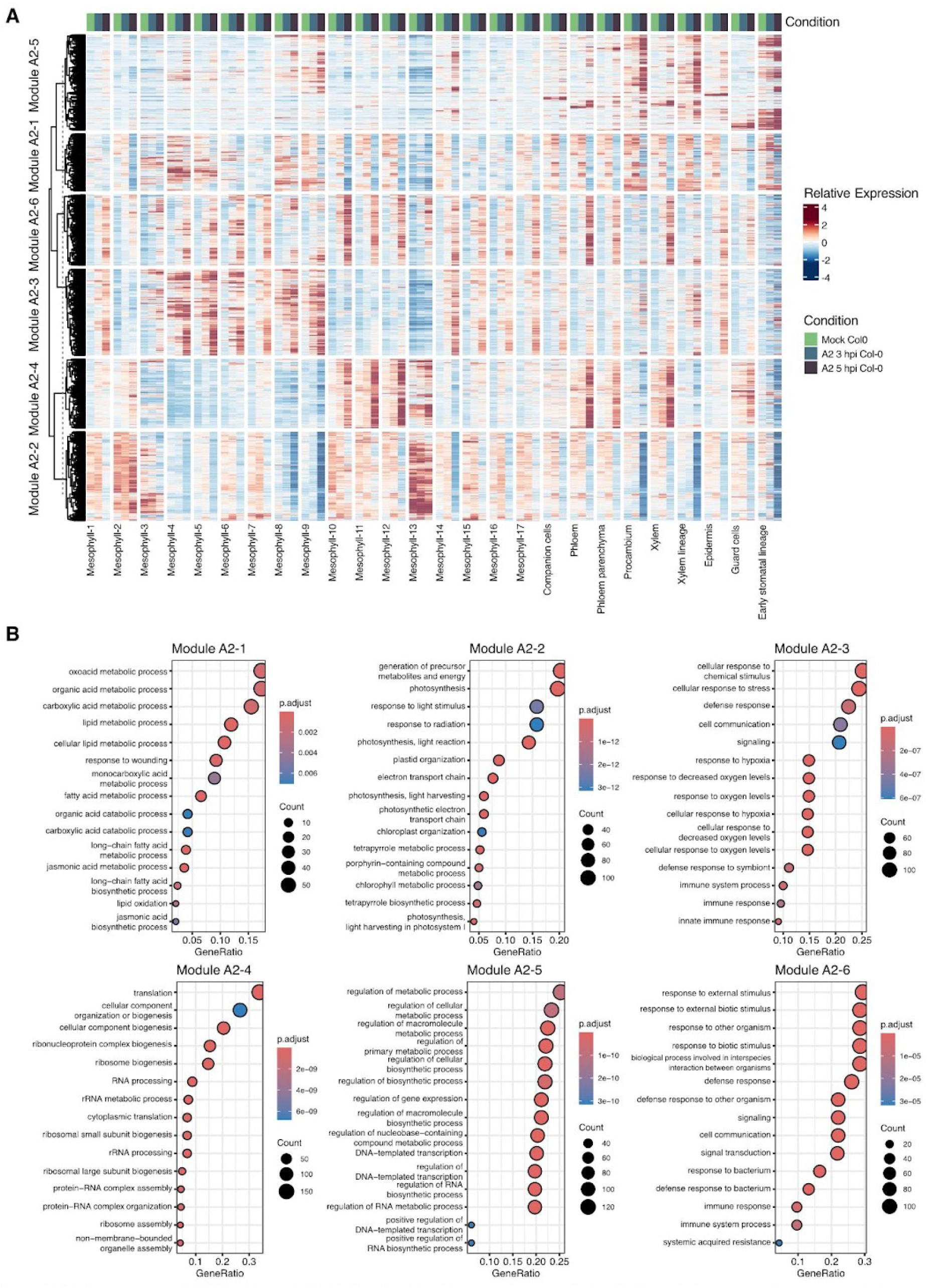
Gene modules in *Pst* DC3000 (AvrRpt2) infected Col-0 samples. **A.** Heatmap of DEGs from all cell clusters of A2 3 hpi/5 hpi vs. mock. All the DEGs were clustered into six co-expression modules (Modules A2-1 to A2-6) based on their expression pattern (k-mean cluster). Columns correspond to the cell clusters; the color bar above indicates the samples: mock (green), A2 3 hpi (light teal) and A2 5 hpi (dark teal). Colour shows the realative expression value (red = high, blue = low). **B.** Functional enrichment of each module. Dotplots show the top enriched GO terms per module (ClusterProfiler, Benjamini–Hochberg FDR). Dot size represents the number of module genes in the term, and color encodes the adjusted *p* value. A2 is short for *Pst* DC3000 (AvrRpt2).

**Fig. S7.**
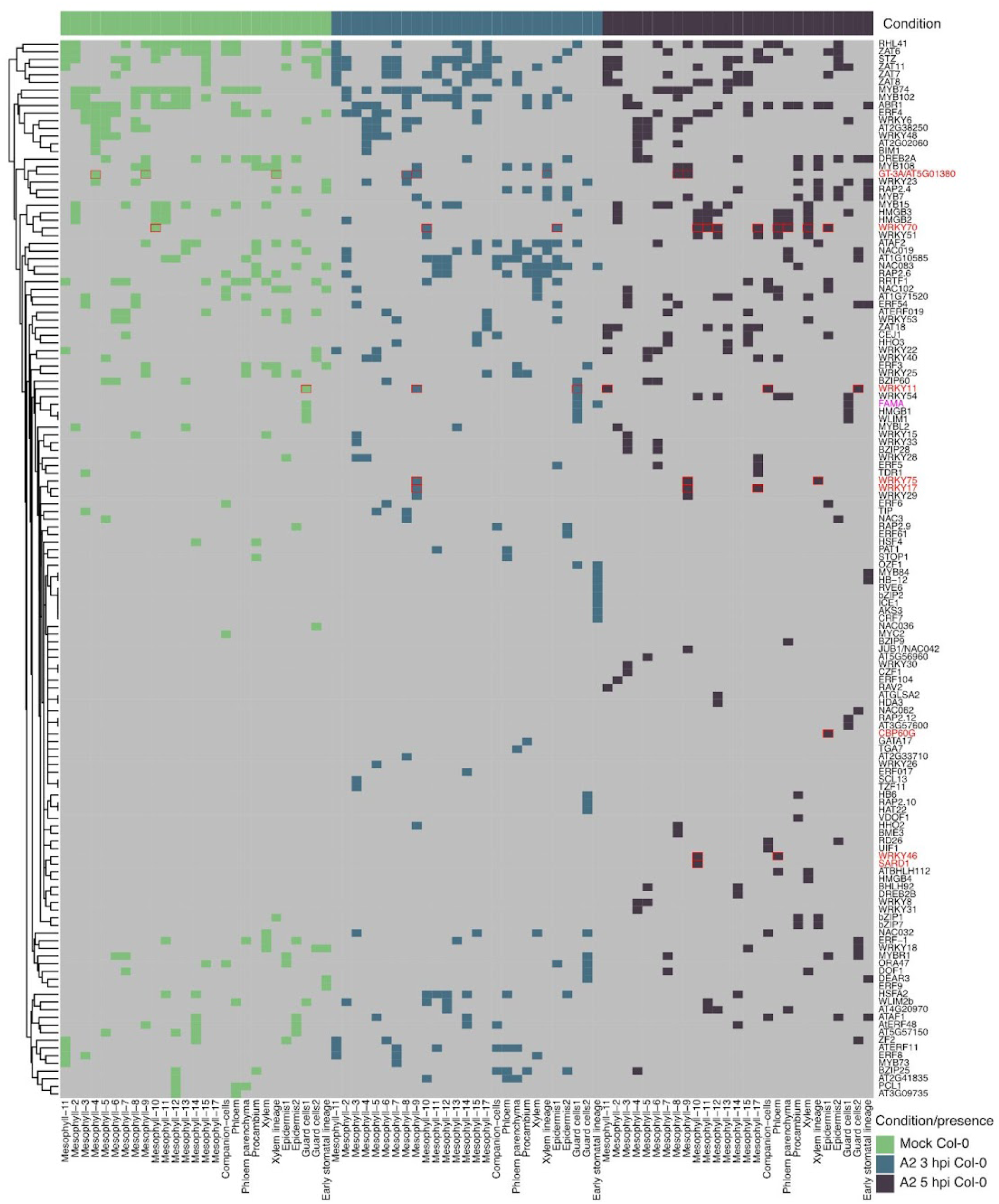
Distribution of the top 10 regulons across cell clusters in *Pst* DC3000 (AvrRpt2) infected Col-0 samples. Presence/absence heatmap summarising the ten highest-ranking specific regulons detected in each cell cluster. The color above indicates the sample: mock (green), A2 3 hpi (light teal) and A2 5 hpi (dark teal). FAMA, GT-3A and SA-related regulons are highlighted.

**Fig. S8.**
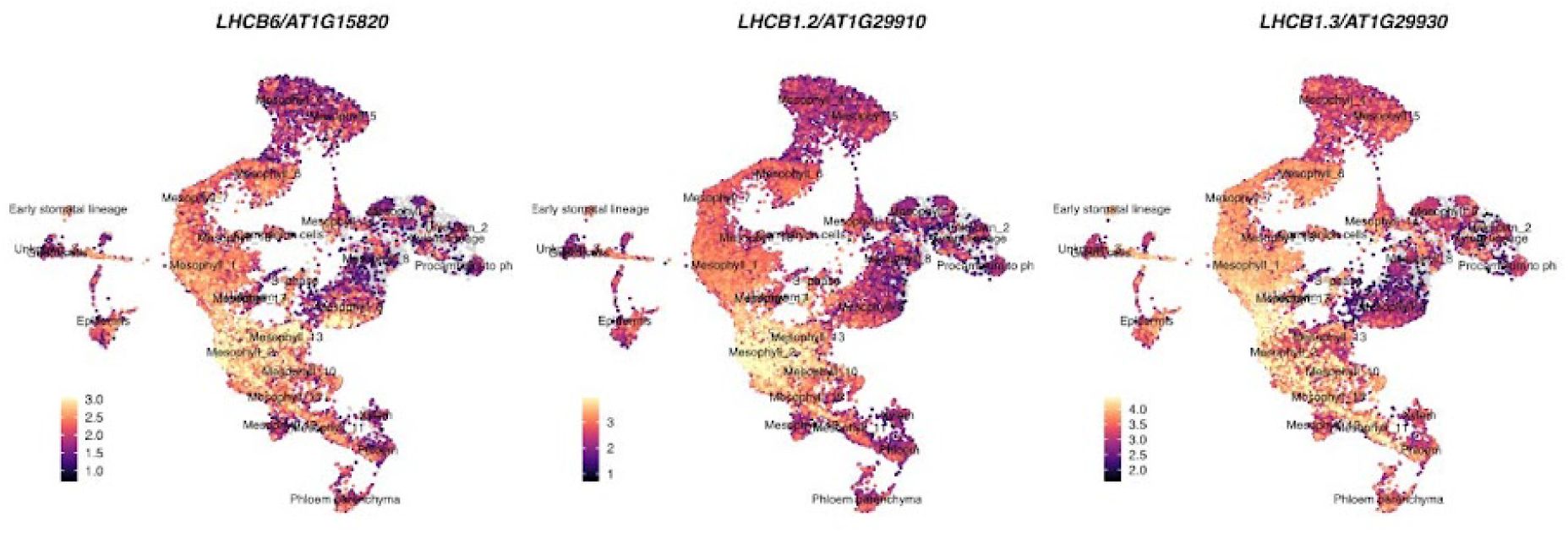
Expression pattern of photosynthesis-related genes. UMAPs of the mock Col-0 atlas colored by scaled expression levels of *LHCB6*, *LHCB1.2*, and *LHCB1.3*. Yellow marks cells with higher transcript abundance, purple marks lower abundance.

**Fig. S9.**
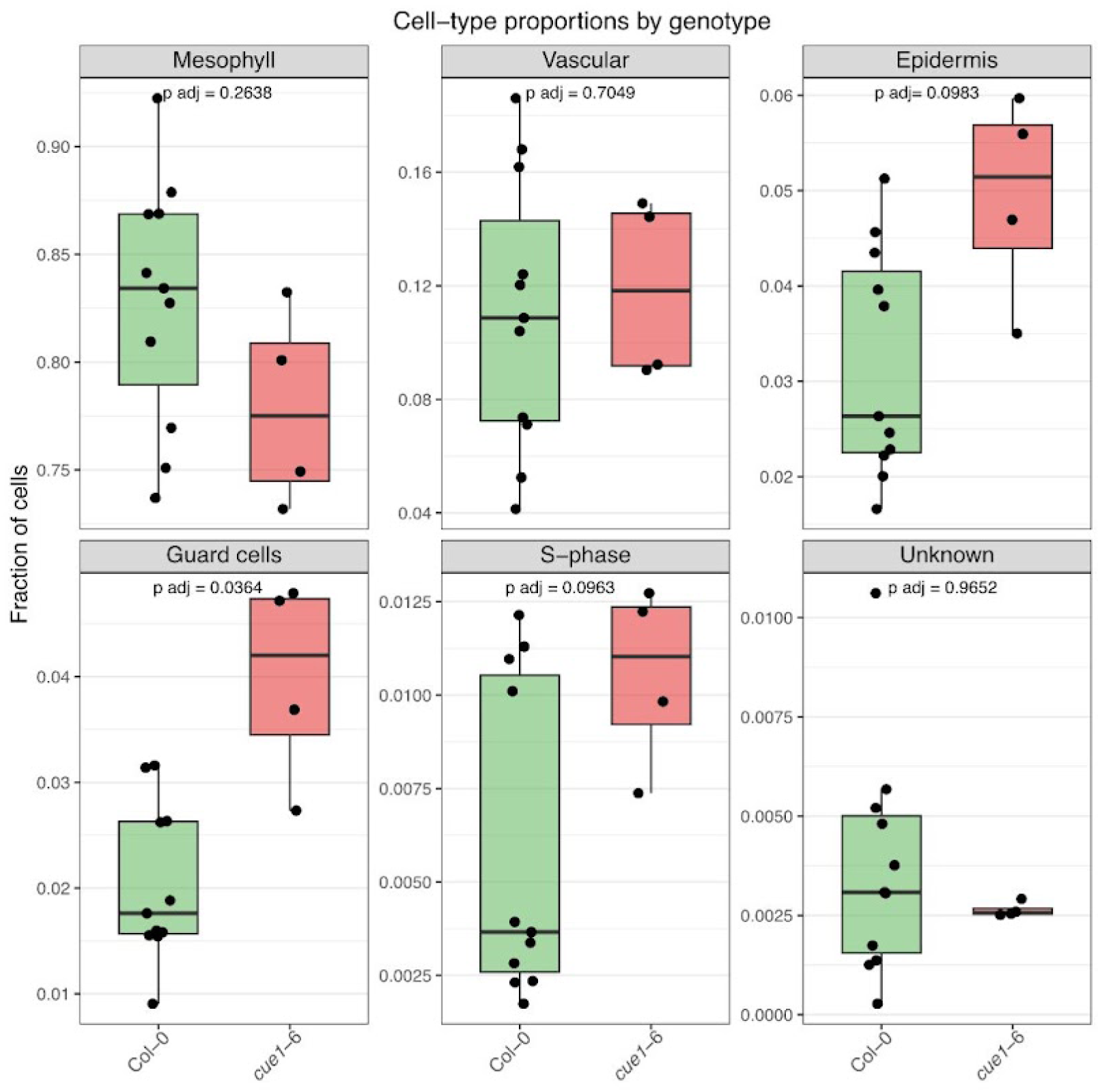
Cell-type proportions in Col-0 versus *cue1*-6. Box plots show the fraction of every major cell type, each dot representing an individual library with Col-0 (n = 11) and *cue1*-6 (n = 4). Benjamini–Hochberg FDR for the moderated *t*-test was performed and the adjusted *p* value is labeled.

**Fig. S10.**
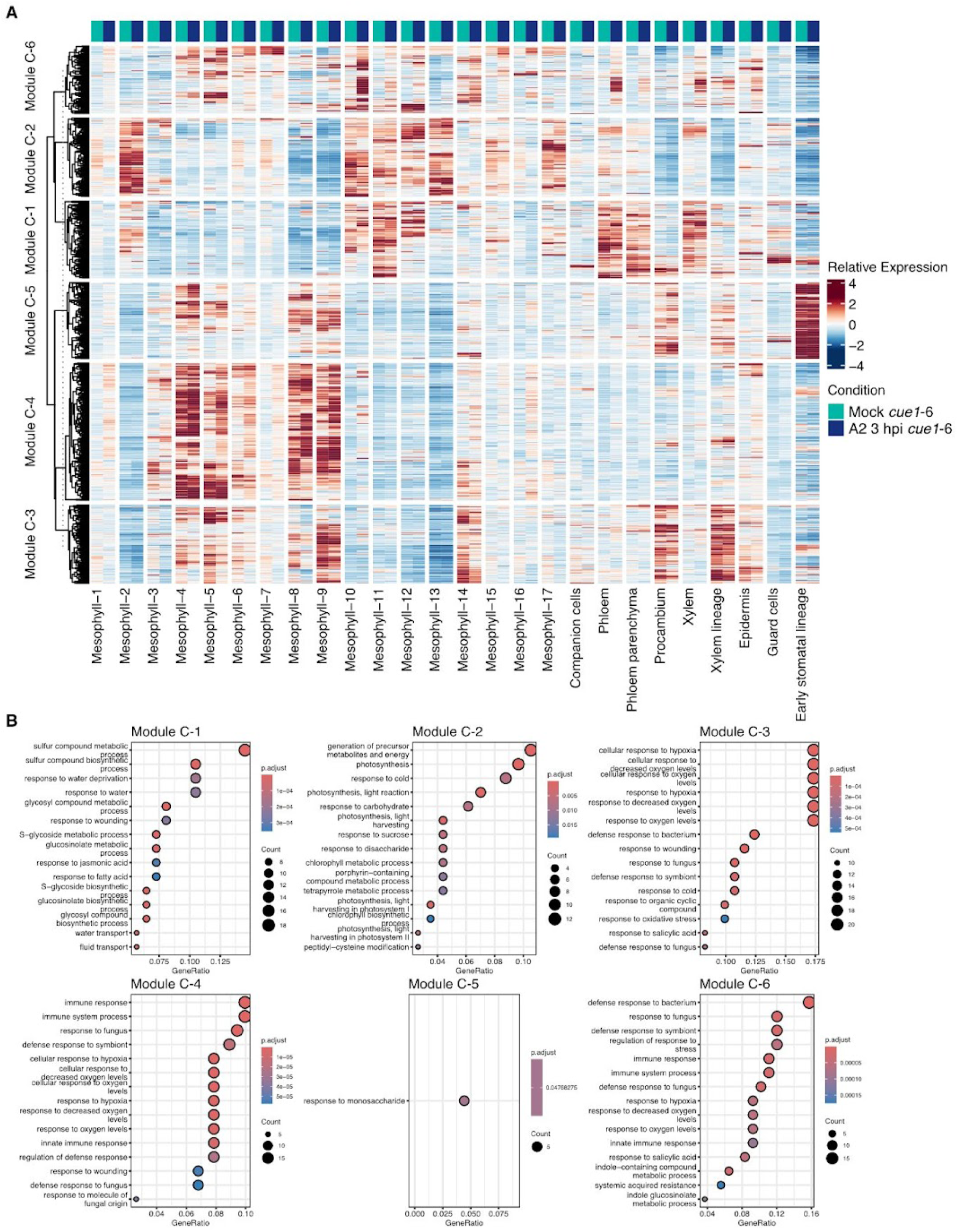
Gene modules in *Pst* DC3000 (AvrRpt2) infected *cue1*-6 samples. **A.** Heatmap of DEGs from all cell clusters of A2 3 hpi vs. mock in *cue1*-6. All the DEGs were clustered into six co-expression modules (Modules *<u>c</u>ue1* (C)-1 to C-6) based on their expression pattern (k-mean cluster). Columns correspond to the cell clusters; the color bar above indicates the samples: *cue1-*6 mock (green) and *cue1-*6 A2 3 hpi (blue). Colour shows the relative expression value (red = high, blue = low). **B.** Functional enrichment of each module. Dotplots show the top enriched GO terms per module (ClusterProfiler, Benjamini–Hochberg FDR). Dot size represents the number of module genes in the term, and color encodes the adjusted *p* value. A2 is short for *Pst* DC3000 (AvrRpt2).

**Fig. S11.**
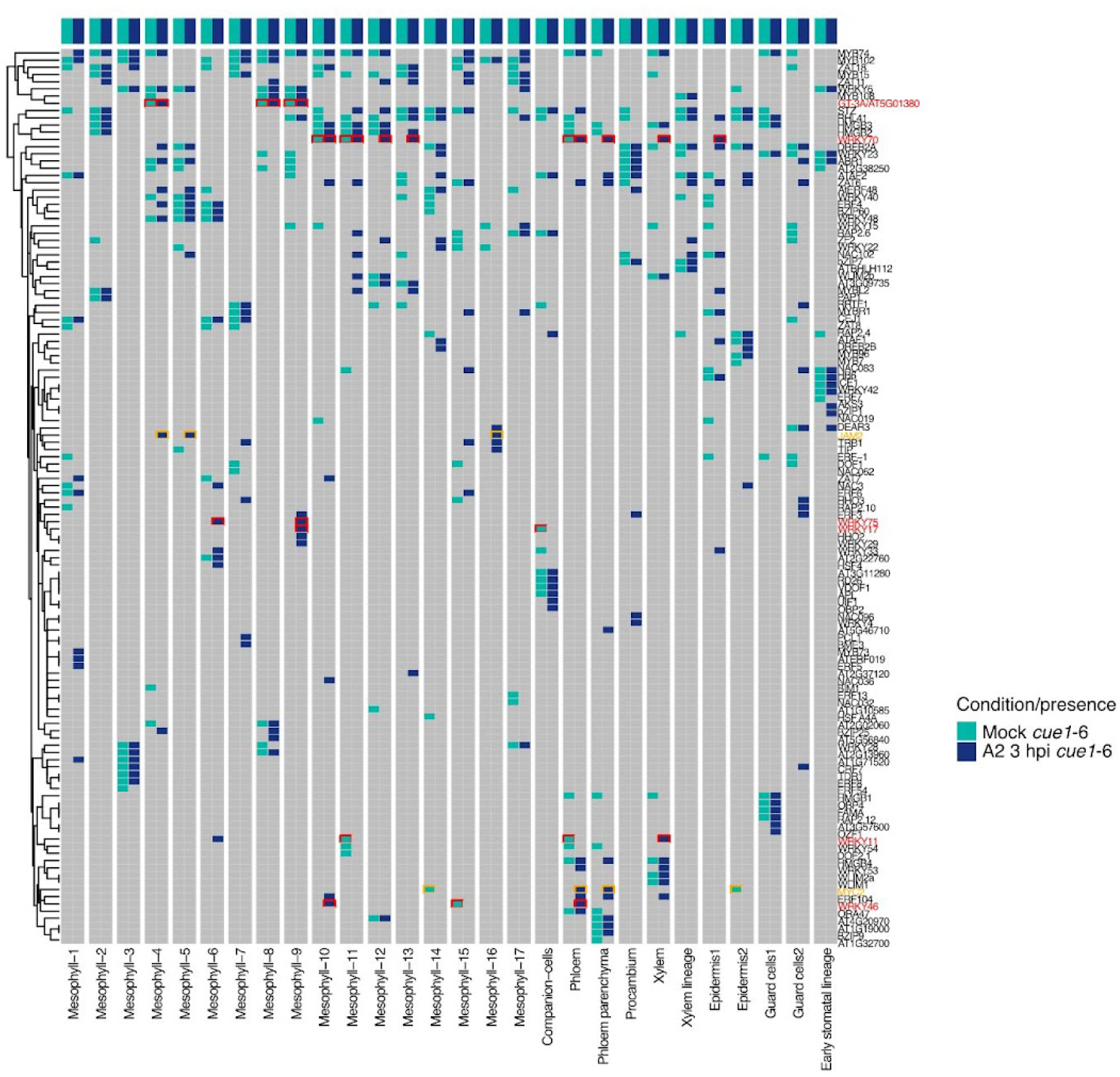
Distribution of the top 10 regulons across cell clusters in *Pst* DC3000 (AvrRpt2) infected *cue1*-6 samples. Presence/absence heatmap summarising the ten highest-ranking specific regulons detected in each cell cluster. The color bar above indicates the sample: *cue1*-6 mock (green) and *cue1*-6 A2 3 hpi (blue). GT-3A and SA-related regulons are highlighted in red, and a JA-related regulon is highlighted in yellow.

### Supplementary Tables

Table S1 | Cell cluster annotation with known cell-type makers.

Table S2 | DEG list from bulk RNA-seq of protoplasts from Arabidopsis leaves infected with *Pst* DC3000.

Table S3 | DEGs from each cell cluster between *Pst* DC3000 and mock treatments in Col-0, and DEGs between *Pst* DC3000 (AvrRpt2) and mock treatments in *cue1-*6.

Table S4 | Gene list from Module EV and Module A2 in Col-0 and Module C in *cue1-*6..

Table S5 | Regulons in each cell cluster from individual samples.

Table S6 | RLK genes and NLR genes examined in this study.

Table S7 | Average expression of RLK genes in *Pst* DC3000 (EV) and NLR genes in *Pst* DC3000 (AvrRpt2) infected samples in Col-0.

